# Triple dissociation of visual, auditory and motor processing in primary visual cortex

**DOI:** 10.1101/2022.06.29.498156

**Authors:** Matthijs N. Oude Lohuis, Pietro Marchesi, Umberto Olcese, Cyriel Pennartz

## Abstract

Primary sensory cortices respond to crossmodal stimuli, for example auditory responses are found in primary visual cortex (V1). However, it remains unclear whether these responses reflect sensory inputs or behavioural modulation through sound-evoked body movement. We address this controversy by showing that sound-evoked activity in V1 of awake mice can be dissociated into auditory and behavioural components with distinct spatiotemporal profiles. The auditory component began at ∼27 ms, was found in superficial and deep layers and originated from auditory cortex, as shown by inactivation by muscimol. Sound-evoked orofacial movements correlated with V1 neural activity starting at ∼80-100 ms and explained auditory frequency-tuning. Visual, auditory and motor activity were expressed by segregated neuronal populations and during simultaneous audiovisual stimulation, visual representations remained dissociable from auditory and motor-related activity. This threefold dissociability of auditory, motor and visual processing is central to understanding how distinct inputs to visual cortex interact to support vision.

## Introduction

During our everyday lives, we sample the world through active exploration with our different senses. Already in primary sensory cortices, contextual signals about ongoing events in other sensory and motor modalities are integrated with modality-specific signals to enable meaningful sensory processing (Ghazanfar and Schroeder, 2006; Guitchounts et al., 2020; Jones and Powell, 1970; Kayser and Logothetis, 2007; Meijer et al., 2019; Niell and Stryker, 2010; Stringer et al., 2019). In general, integration of crossmodal and motor information may enhance sensory detection and discrimination by way of Bayesian cue integration, may subserve cross-modal predictions, and may underlie the contextual and modally distinct representations characteristic of conscious experience (Fetsch et al., 2013; Pennartz, 2015, 2009). In rodents, primary and secondary visual and auditory cortices share direct anatomical connections (Budinger et al., 2006; Budinger and Scheich, 2009; Campi et al., 2010; Cappe and Barone, 2005; Falchier et al., 2002; Miller and Vogt, 1984; Paperna and Malach, 1991; Rockland and Ojima, 2003) and auditory inputs to primary visual cortex (V1) have been found to target L1 and L5/L6 (Ibrahim et al., 2016; Iurilli et al., 2012; Mesik et al., 2019; Rockland and Ojima, 2003). Auditory inputs affect visual response properties such as orientation and contrast tuning in V1 (Ibrahim et al., 2016; Meijer et al., 2017), with some V1 neurons directly responding to sounds (Meijer et al., 2017). Even more strikingly, selectivity to auditory features such as spatial location and frequency has been reported in cat visual cortex (Fishman and Michael, 1973; Morrell, 1972; Spinelli et al., 1968) and mouse V1 (Knöpfel et al., 2019).

In addition to crossmodal inputs, other factors strongly impact activity in primary sensory areas. Locomotion and increases in arousal both desynchronize spike patterns and increase V1 activity (Fu et al., 2014; Niell and Stryker, 2010; Vinck et al., 2015). Also eye, head and orofacial movements lead to marked activity changes (Bouvier et al., 2020; Guitchounts et al., 2020; Stringer et al., 2019). The exact function of motor signaling to primary sensory cortices remains unknown, but it possibly provides an efference copy serving to predict the sensory consequences of body movement. Failing to account for these motor-related influences risks misinterpreting the observed sensory cortical activity (Musall et al., 2019; Zagha et al., 2022). Indeed, V1 activity during sound clips has been argued to have a behavioral rather than sensory origin and relate to stereotyped sound-evoked movements (Bimbard et al., 2021). The origins of crossmodal activity in sensory cortex are thus unclear and dissociating auditory from behavioral signals in V1 is central to correctly interpret (multi-)sensory activity and to understand how distinct inputs to visual cortex interact to support vision.

To examine whether visual, auditory and motor processing can be dissociated in V1, we trained mice on a task which required them not only to detect sensory changes, but also to distinguish or segregate the modality in which the change occurred. In awake animals performing this task, auditory stimuli evoke frequency-tuned neuronal responses in V1, that are largely, but not completely, explained by orofacial movements. We disentangled auditory-related from motor-related activity using multi-area recordings, task manipulations, pharmacological interventions and optogenetics. An early, sound-evoked component of V1 responses originates from auditory cortex, is transient, and is found predominantly in superficial and deep layers. In contrast, motor-related activity following auditory stimuli results from rapid orofacial movements with an onset-latency of 60-100 ms, is found mostly in superficial layers, and underlies the bulk of late sound-evoked neural activity changes in visual cortex, but not in auditory cortex. Jointly, these signals strongly affected visual cortical activity in a way that leaves visual stimulus coding intact.

## Results

To disentangle visual, auditory and behavior-related signaling in V1 we presented three different cohorts of head-fixed mice with the same sensory stimuli, but trained them to report only visual stimuli, both auditory and visual stimuli, or neither (Fig. 1a). Visual stimuli were continuous full-field drifting gratings and visual trials consisted of uncued, occasional orientation changes. Auditory stimuli consisted of weighted combinations of five harmonic tones that occasionally changed frequency (auditory trial; Ext. Data Fig. 1a-c). Noncontingently exposed mice (NE; N=7) were not rewarded for licking after any stimulus change, but were pseudorandomly rewarded for spontaneous licks and therefore served as naive control animals. Unisensory trained mice (UST; N=4) were only rewarded for lick responses to visual changes and learned to ignore auditory changes. In the multisensory task version (MST; N=17), responses to both visual and auditory changes were rewarded and animals were trained to lick left to auditory changes and right to visual changes (or vice versa). Thus, not only the change but also modality identity was behaviorally reported through directional lick responses. Hit rate scaled with stimulus saliency (amount of visual or auditory feature change) only in modalities for which mice were rewarded (Fig. 1b), confirming effective manipulation of the task-relevance of sensory changes.

**Figure 1:**
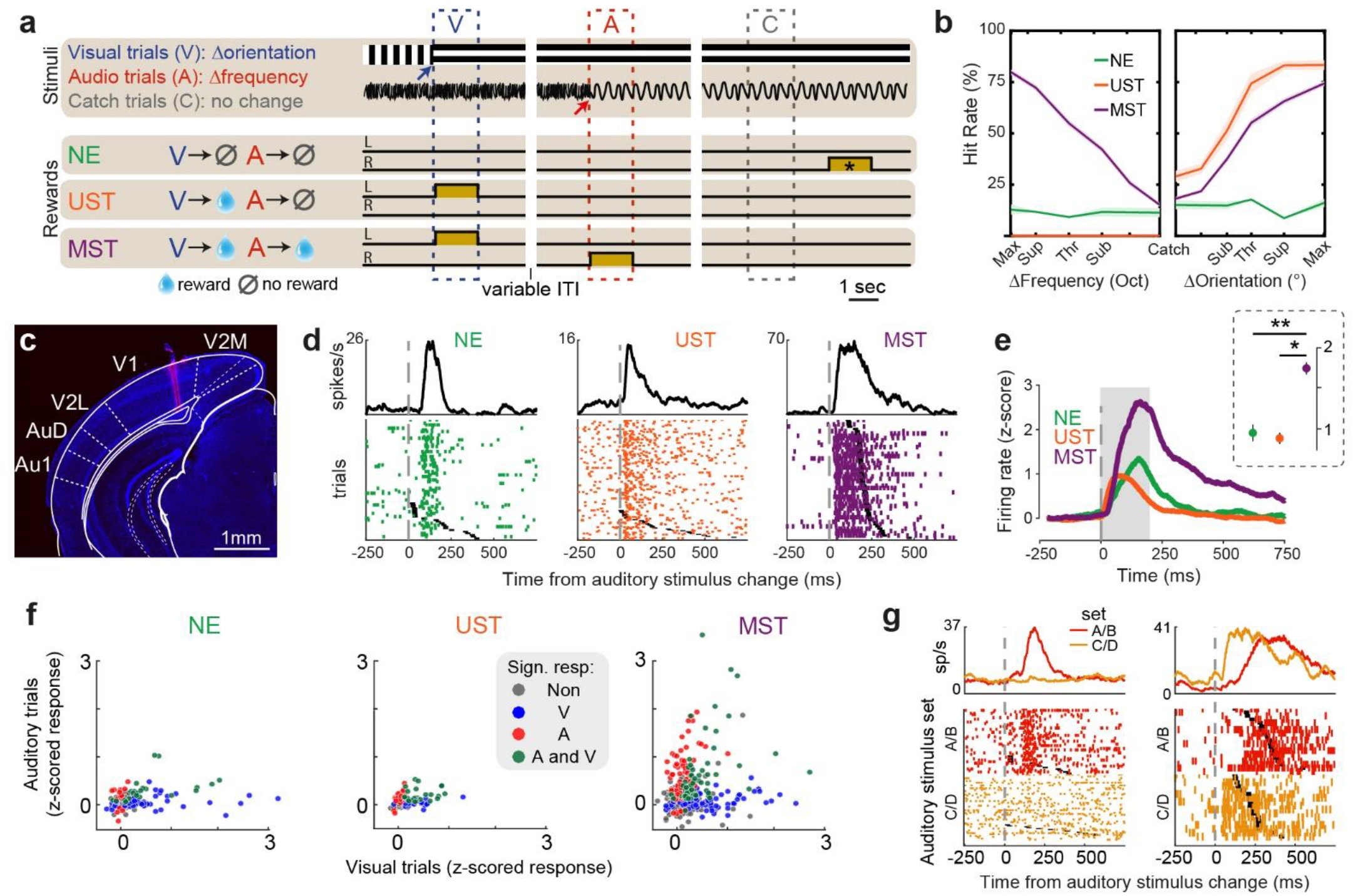
Task dependent recruitment of auditory activity in visual cortex upon behavioral relevance. **a)** Stimuli and reward contingencies for the three cohorts of mice. Visual and auditory stimuli were continuously presented and visual trials (V) consisted of uncued orientation changes (blue arrow), auditory trials (A) of uncued frequency changes (red arrow), and catch trials (C) of no change. Noncontingently exposed mice (NE) were not rewarded for licking after stimulus changes, unisensory trained mice (UST) for licks after visual but not auditory changes, and multisensory trained mice (MST) for reporting both visual and auditory changes. In MST mice, each lick spout was associated with a single modality. Ochre rectangle indicates reward availability. *For NE mice, reward windows were temporally decorrelated from the sensory stimuli, and randomly occurring outside the stimulation period. The dotted trial windows indicate the time window used post-hoc to compare stimulus-related lick rates across cohorts. ITI: inter-trial interval. **b)** Average hit rates at different stimulus saliencies for each cohort. Sub: subthreshold, thr: threshold, sup: suprathreshold, max: maximal). Behavioral hit rates increased as a function of the amount of auditory change (step size in frequency) in MST mice (left panel), and as a function of visual orientation change for UST and MST mice (right panel), in line with their reward schedule. UST mice were only rewarded for licks to the visual lick spout and therefore chose not to lick the auditory lick spout. Mean ± SEM. **c)** DAPI-stained (blue) coronal section showing electrode track in left V1 stained with DiI (red) ±3.56 mm posterior to Bregma. V2M and V2L: medial and lateral secondary visual cortex, respectively. Au1: primary auditory cortex. AuD: dorsal secondary auditory cortex. **d)** Raster plots showing sound-evoked activity in example V1 neurons from each cohort. Black ticks indicate the first lick after stimulus change, which was rewarded only in MST mice. **e)** Averaged z-scored firing rate (referenced to baseline) for auditory responsive V1 neurons to preferred post-change auditory stimulus. Inset shows response averaged across the shaded analysis window (0-200 ms). *p<0.05, **p<0.01. Mean ± SEM. **f)** Scatter plot of z-scored firing rate, corrected for baseline activity (analysis window 0-200 ms) and following auditory and visual stimulus changes. Each dot is a neuron. (NE: N=163; UST: N=128; MST: N=812). Colors denote cells with a significant response to any modal input (gray: no significant response). **g)** Individual V1 neurons show frequency-specific sound-evoked activity. Raster plots as in (d), but for the two sets of auditory post-change frequencies (where A and B are two similar frequencies and C and D as well; Ext. Data Fig. 1e). Left example from NE mouse, right from MST mouse.

### Sound-evoked activity in primary visual cortex

We recorded single unit and LFP activity in V1 (Fig. 1c). During recording sessions, two levels of change saliency were presented per modality: threshold level (Vthr and Athr, individually titrated per animal based on psychophysical data; see Methods) and maximal saliency (Vmax and Amax, 90° and ½ octave change, respectively; Ext. Data. Fig. 1d). Furthermore, stimuli were restricted to four orientations and four frequencies. Within these visual or auditory stimulus sets, stimuli A and B were highly similar (differing at threshold level), as were C and D, and nearby stimuli were grouped into pairs for analyses (set A/B vs. set C/D; see Ext. Data Fig. 1e).

Auditory frequency changes at maximal saliency (Amax) induced spiking activity in V1 neurons of mice from all three cohorts (Fig. 1d). These responses were generally transient and subsided after 200 ms. A sizeable fraction of V1 neurons had a significantly higher firing rate (0-200 ms) compared to baseline following at least one of the post-change auditory stimuli (Wilcoxon signed rank test, p<0.05; during Amax trials). This fraction was similar across cohorts (NE: 25.5 ± 4.0%, n=5 sessions; UST: 34.8 ± 4.8% n=4 sessions; MST: 20.9 ± 0.7% n=27 sessions; p=0.30, Kruskal-Wallis test). A large fraction of V1 neurons also responded to one of the two visual orientation changes in all three cohorts (NE: 42.4 ± 7.9%; UST: 53.2 ± 6.8 %; MST: 30.2 ± 0.8%; mean ± SEM; p=0.13, same sessions as above, Kruskal-Wallis test). To quantify and compare neuronal activity, firing rates during auditory and visual trials were z-scored (Fig. 1e,f). Auditory responses of V1 neurons (0-200 ms after stimulus change) were particularly strong in MST animals, in which responses exceeded those observed in NE and UST animals (F(2,232)=4.96, p=0.01; Posthoc comparison: MST vs NE: F(1,229)=4.40, p=0.037; NE vs UST: F(1,229)=0.02, p=0.876; MST vs UST: F(1,229)=7.75, p=0.006). V1 neurons were previously found to be selective for auditory stimulus features such as sound frequency (Fishman and Michael, 1973; Knöpfel et al., 2019; Morrell, 1972; Spinelli et al., 1968). Indeed, individual neurons displayed stimulus-specific sound-evoked firing rate responses (Fig. 1g) and a surprisingly large fraction of V1 neurons discriminated post-change auditory stimulus identity (12.7% ± 3.7%, Mean +-SEM across sessions, A/B vs C/D Amax trials, permutation test; p<0.05), which was comparable across cohorts (NE: 11.9 ± 2.7%; UST: 16.1 ± 4.5%; MST: 12.1 ± 1.9%; p=0.68, Kruskal-Wallis test). This was significantly smaller than the fraction of orientation-selective cells across cohorts (31.9 ± 3.7%; fraction frequency-versus orientation-tuned: F(1,52)=18.48, p=7.54*10^−5^). This fraction is lower than commonly reported in the literature, presumably because we only sampled four irregularly spaced grating orientations (Ext. Data Fig. 1e).

### Motor contribution to sound-evoked activity in visual cortex

The enhanced sound-evoked responses in V1 of MST animals are potentially explained by licking as only these mice were rewarded for reporting auditory changes. Auditory lick reaction times in MST mice were around 320 ms (median, IQR: 219-446 ms) and faster than visual reaction times (median: 404 ms; IQR: 320-509 ms; F(1,270) = 105.65, p=3.94 * 10^−21^), suggesting motor-related activity changes in V1 could already play a role during early time windows after the change in sound (0-200 ms). We recorded the mice’s faces and often observed orofacial movements following auditory stimulus changes in trained, but also untrained mice. We therefore explored to what extent sound-evoked activity in V1 was related to (stereotypical) movements (Bimbard et al., 2021; Williams et al., 2021). To investigate this, motion energy was extracted from video footage (video ME) as the overall pixel intensity difference between consecutive frames (Fig. 2a), which included snout, whisking and licking movements.

**Figure 2.**
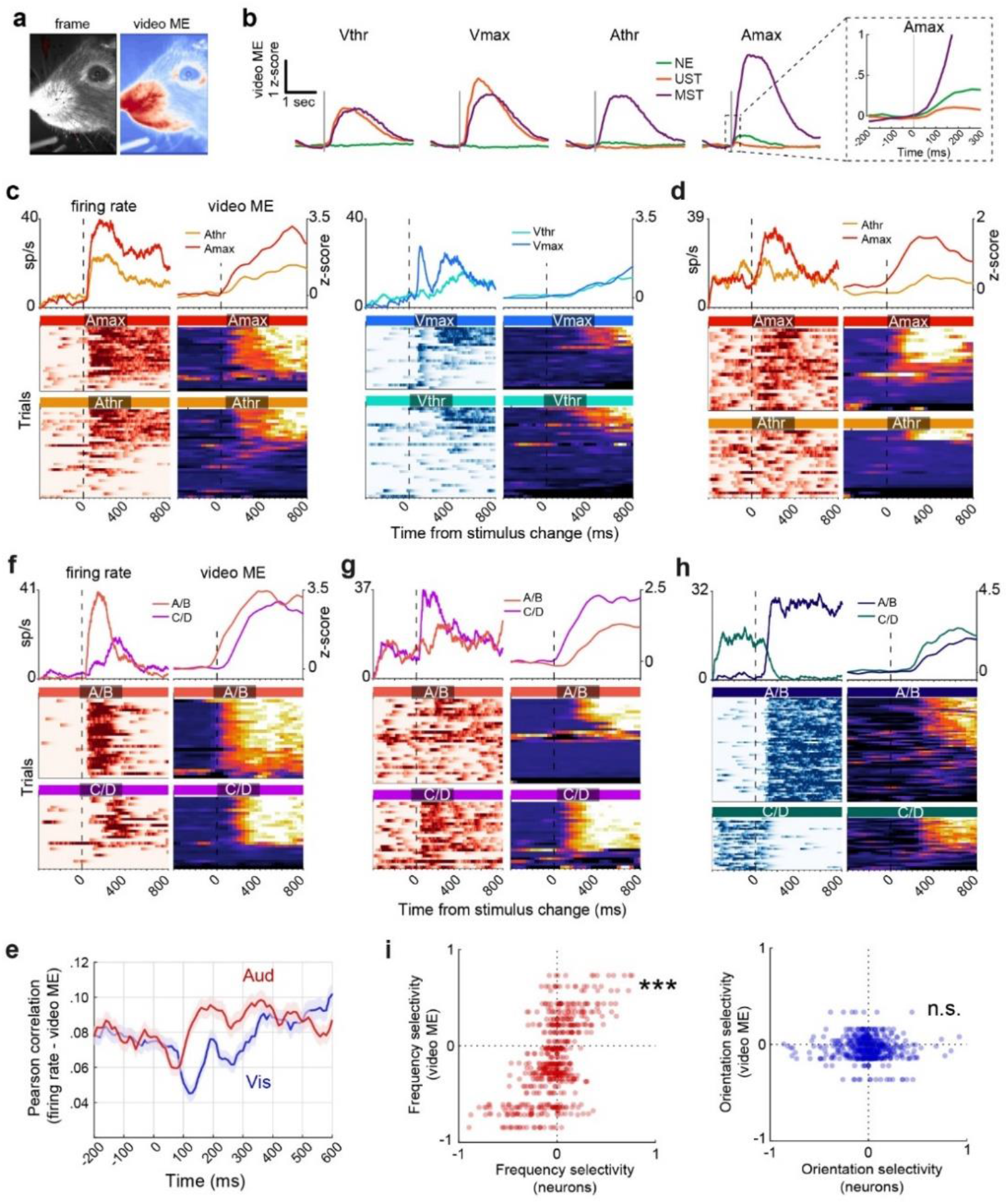
Sound-evoked orofacial movements explain frequency-tuned activity in visual cortex. **a)** Example frame and motion energy from the video (video ME). Red indicates more orofacial movements. **b)** Video ME for auditory (threshold and maximal saliency, Athr and Amax) and visual (Vthr and Vmax) trials across cohorts. Inset shows a zoom in of boxed region in Amax trials. **c)** Trial-by-trial spiking activity correlates with video ME in an example V1 neuron (MST animal). Columns from left to right: firing rate during auditory trials (red is high firing rate), video ME during auditory trials (warm colors is increased orofacial motion), firing rate during visual trials (blue is high firing rate), video ME during visual trials. Trials include hits and misses and are separated by saliency (thr and max) and sorted by magnitude of video ME. Upper panels show trial-averages of heatmaps below. Heatmap color range is the axis range of the upper plot. **d)** Same as c, but for another example V1 neuron (MST cohort) with sound-evoked spiking activity in Amax trials without changes in video ME in a subset of trials. **e)** Average Pearson correlation of single neuron firing rate to video ME during auditory (red) and visual (blue) trials over time. **f-h)** Same as *c*, but with trials separated based on visual and auditory post-change stimulus class (set A/B versus set C/D). In the left two examples, both firing rate and video ME discriminate post-change auditory feature, and firing rate and video ME are correlated across trials. In the right example, firing rate discriminated grating orientation independently of orofacial movements and video ME was largely unrelated to grating orientation. Pre-change coding resulted from the fact that gratings changed from A/B to C/D, or vice versa. Neurons from MST mice. **i)** Left: frequency selectivity of V1 neurons (x-axis) and video ME-based selectivity in the same session (y-axis). Selectivity was measured as AUC, rescaled between -1 and 1. Each dot is a neuron (all cohorts combined). Right: same but for orientation selectivity.

Visual stimulus changes induced movements only in UST and MST mice, the cohorts rewarded for licking. A change in auditory stimulus was followed by strong movements in MST mice, but to a lesser extent also in NE and UST mice (Fig. 2b). In Amax auditory trials, these orofacial movements started roughly 60-100 ms after the stimulus change, but our temporal resolution was limited by a frame rate of 25 fps. In MST animals, instrumental licking movements dominated, while in NE and UST animals, where sounds were behaviorally irrelevant, they evoked mainly whisking and snout movements (Ext. Data Fig. 2a,b). Similarly, if we removed the lick spout in MST mice, auditory stimuli continued to evoke orofacial movements other than licking, while movements disappeared altogether following visual stimuli (Ext. Data Fig. 2c-e). Sounds thus evoked orofacial movements whether relevant or irrelevant to the current task, in contrast to visual stimuli, which only led to reward-contingent movements.

### Sound-evoked movements underlie frequency tuning

For of a number of individual V1 neurons, activity correlated with video ME (Ext. Data Fig. 3a) (Stringer et al., 2019). Also during auditory trials, some neurons showed an overall similarity between trial-to-trial firing rate and video ME (Fig. 2c). However, example neurons also showed sound-evoked activity in trials without increases in video ME (Fig. 2d). The increase in orofacial movements after auditory changes could therefore underlie sound-evoked activity in V1 at least in a fraction of neurons. As a first test of this hypothesis, we correlated firing rate with video ME during auditory and visual trials for all neurons (Fig. 2e). Relative to baseline, the average correlation decreased for visual trials, likely due to visually driven activity unrelated to ongoing movements. Following auditory changes, the average correlation transiently dropped and then increased after roughly 100 ms, in line with the observed movement-related activity of individual V1 neurons.

We next wondered whether movements could also underlie the observed auditory frequency-specific V1 activity (Fig. 1g). As illustrated for two example neurons (Fig. 2f,g), frequency-tuned activity was strongly aligned to variability in video ME in those trials. Both neuronal activity and video ME thus responded to auditory changes in a stimulus-specific manner. This was not the case for orientation-tuning (Fig. 2h). Auditory feature tuning could therefore result from ‘motor-tuning’. Indeed, across the full population, tuning to either one of the auditory stimuli (quantified using the receiver operating characteristic, ROC) was accompanied by strong movements to that stimulus (F(1,688)=286.27, R=0.542, p=5.92*10^−54^), but this was not the case for visual neuronal selectivity (F(1,692)=2.90, R=0.065, p=0.09; Fig. 2i). Furthermore, if strong motor activity drove frequency selectivity, one would expect all simultaneously recorded cells to be preferentially tuned to the same auditory stimulus (viz. the one that elicited most movement). Indeed, simultaneously recorded V1 neurons responded to the same auditory stimuli, while their preferred grating orientation was mixed (Ext. Data Fig. 3b). These findings were corroborated in a separate set of animals trained on a stimulus detection (rather than change detection) version of the task, which allowed us to test a more extended range of visual and auditory stimuli (see Methods; Ext. Data Fig. 4). We found similar results when training population decoders on V1 population spiking activity or on high-dimensional video data (Ext. Data Fig. 3c-k; dimensions are understood here as different PCA components; Stringer et al. 2019).

Auditory frequency could be decoded from neural and video data in a highly correlated manner, whereas grating orientation could be deduced only from V1 activity. In sum, auditory stimuli led to fast orofacial movements that correlated with sound-induced and tone-specific V1 activity.

### Dissociable auditory and motor-related signals

A purely behavioral origin of sound-evoked activity in V1 would be in disagreement with some previous findings. Primary auditory and visual cortices are monosynaptically connected and sound-evoked activity has been reported under anesthetized conditions where overt movements and changes in behavioral state play a very minor role, if any (Henschke et al., 2015; Ibrahim et al., 2016; Iurilli et al., 2012). Furthermore, in our recordings, many neurons showed short-latency sound-evoked activity also in trials without apparent changes in motor activity (e.g. Fig. 2d). We therefore explored whether auditory activity can be disentangled from motor-related activity in visual cortex.

First, we compared sound-evoked V1 activity to spiking activity in auditory cortex, where the importance of bottom-up auditory signaling is well established (AC; including primary auditory cortex, anterior auditory field and dorsoposterior auditory cortex; Fig. 3a). AC strongly responded to auditory stimulus changes, but only minimally to visual orientation changes (Fig. 3b, *right*). We aggregated population activity across neurons to achieve a higher temporal resolution than single-unit activity and compared the latency to firing-rate increases evoked by visual and auditory stimuli in V1 and AC (Fig. 3b). The auditory response started at 18 ms in AC and 27 ms in V1. The small increase in AC activity after visual stimuli became significant only after 156 ms. This short onset latency of sound-evoked activity in visual cortex fits auditory-related inputs to V1. Visual responses started at 54 ms in V1, matching the canonical retinogeniculate drive (Schnabel et al., 2018).

**Figure 3.**
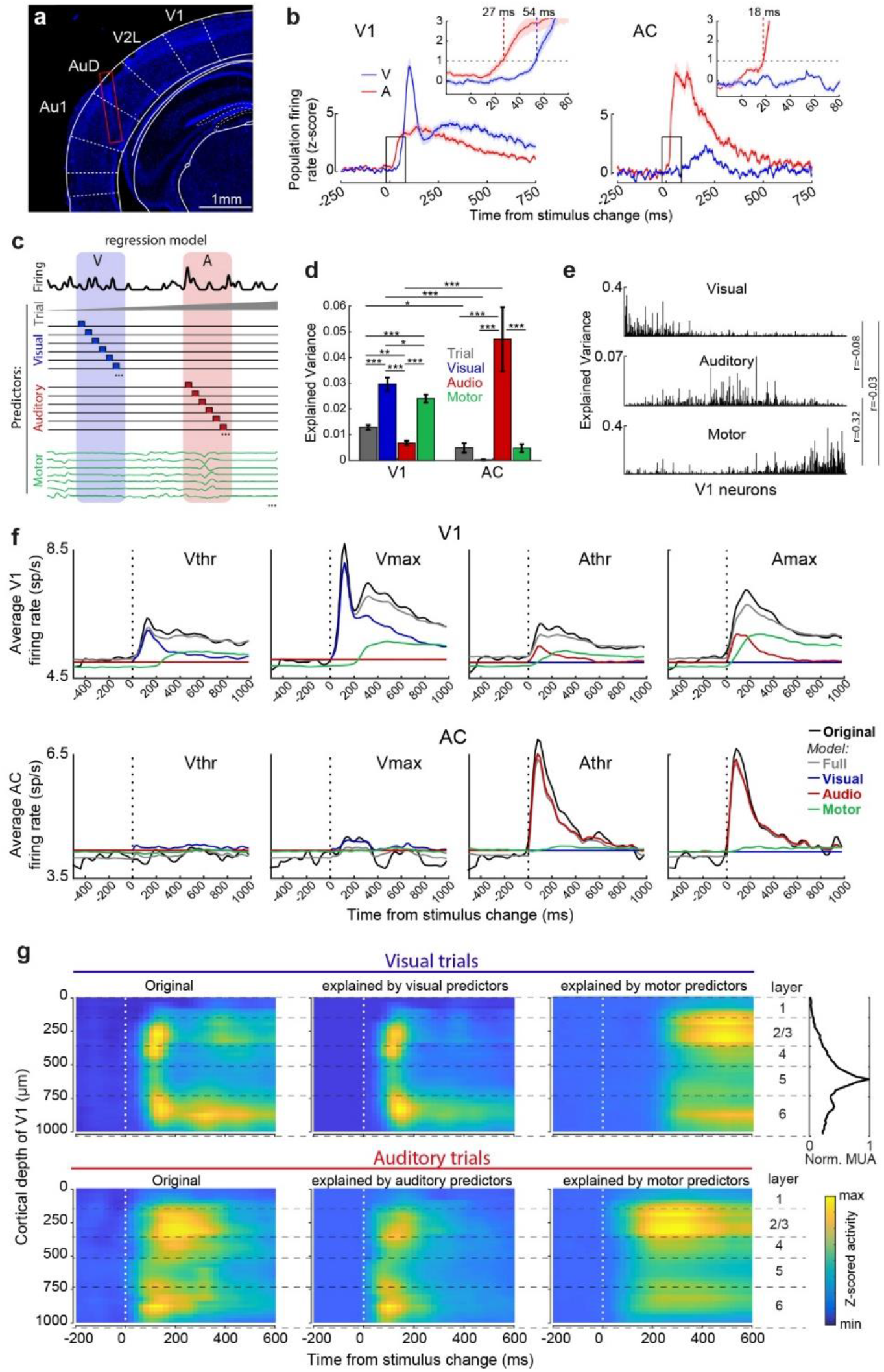
Temporal and spatial dissociation of auditory and motor-related activity in auditory and visual cortex. **a)** DAPI-stained (blue) coronal section showing electrode track in Auditory dorsal (AuD) and primary Auditory cortex (Au1) approximately ±2.46 mm posterior to Bregma. **b)** Population firing rate after auditory and visual stimulus changes in V1 (left) and AC (right). Inset shows close-up of boxed region and the latency to cross a threshold of Z-score > 1. Line and shaded region are mean and SEM across sessions. **c)** Schematic of regression model. Single neuron firing rate was predicted as a linear combination of four sets of predictors. A ‘Trial’ predictor captured rate fluctuation throughout the session. For visual (V) and auditory (A) stimuli a set of separate predictors spanned one second after stimulus change per saliency level and feature (orientation and auditory frequency; shown only for one trial type) to capture neuron-specific response patterns. Motor predictors included the top 25 video PCs. **d)** Explained variance during 0-200 ms of all trials for each category of predictors. Mean ± SEM across neurons. Results based on 51 sessions; NE: 9, UST: 10, MST: 32 sessions, 19.451 trials, 790 V1 and 99 AC neurons. Asterisks indicate significance of a post hoc F-test on the relevant contrast in the linear mixed effects model. *p<0.05, **p<0.01, ***p<0.001. **e)** Distribution of explained variance across V1 neurons for each category of predictors (except Trial), showing encoding of sensory and motor variables in distinct and overlapping neurons. Neurons are sorted based on visual minus motor encoding and centered by auditory encoding. **f)** The firing rate of single neurons was predicted using only subsets of predictors (visual: blue, auditory: red, motor: green) or all predictors (gray) and compared to the original firing rate (black). Shown is the firing rate during different trial types averaged across V1 neurons (upper row) or AC neurons (bottom row). **g)** Heatmap of z-scored firing rate with V1 neurons binned by cortical depth. Leftmost heatmaps show original data; middle and right panels show predictions using sensory or motor variables from the regression model. Rightmost panel shows normalized multi-unit activity peaking in L5 used to demarcate layers (see Ext. Data Fig. 6). Throughout the figure neurons from all cohorts were combined. See Ext. Data Fig. 5b,c for cohort-specific findings.

To further disentangle auditory and motor-related signals, we built a regression model (Fig. 3c) to predict single-neuron activity based on trial number (e.g. to account for drift in motivational state), visual and auditory stimulus features and motor activity (the first 25 video PCs). Given that video predictors represented orofacial movement, we termed the correspondingly explained neural activity *motor-related*.

This approach succeeded in separating the temporally overlapping contributions of sensory stimuli and motor-related activity to trial-by-trial single-neuron firing rate (Ext. Data Fig. 5a). Visual stimuli explained most V1 variance during 0-200 ms post-stimulus change, followed by motor-related activity as next best predictor (Fig. 3d). Auditory stimuli explained a modest but significant fraction of the variance. Motor activity explained roughly three times as much variance as the auditory stimuli. In auditory cortex, firing rate was best explained by auditory stimuli, with only a minor contribution of trial-number and motor activity, and no apparent influence of visual stimuli. In V1, visual encoding was found largely in a separate set of neurons (Fig. 3e) and visual EV was not correlated to motor EV (r=-0.03, F(1,471)=0.61, p=0.44), nor auditory EV (r=-0.08, F(1,484)=3.49, p=0.06). Neurons were observed that uniquely encoded auditory or motor features, as well as jointly auditory-motor coding neurons and the overall EV was moderately correlated (R=0.32, F(1,473)=40.56, p=4.52×10^−10^).

To examine how auditory or motor-related activity distinctly contributed to the average response in V1, single neuron firing rate was predicted using only subsets of predictors in our model (Fig. 3f). Averaging this predicted activity across V1 neurons revealed that activity during visual trials was mostly visual in the early phase (0-200 ms), with a late motor-related component (>200 ms). Activity during auditory trials presented a combination of temporally overlapping auditory and motor-related components, with only early activity (<100 ms) being predominantly auditory. This was strikingly not the case in auditory cortex, which showed a negligible contribution of motor activity to the averaged response. The sound-evoked activity in V1 was thus composed of a distinct, early auditory and a later motor-related component. Performing these analyses for the different cohorts showed that auditory-related activity in V1 was similar across cohorts and that the larger sound-evoked response in MST mice (Fig. 1e) was likely the result of increased motor-related activity due to instrumental licking in this cohort (Ext. Data Fig. 5b,c).

### Laminar organization of auditory and motor-related inputs to V1

As V1 was recorded with microelectrode arrays that spanned the different layers, we next wondered whether the auditory and motor-related components had distinct spatiotemporal profiles. The electrode position was aligned to the cortical depth for each V1 penetration (Ext. Data Fig. 6, Methods). We constructed heatmaps of z-scored firing rate as a function of cortical depth and time relative to the stimulus change, using either the raw data or the activity predicted from stimulus or motor components only (Fig. 3g). Visually induced spiking activity clustered in layer 4 (L4) to L2/3 as well as later in L6, while motor-related activity accounted for the later component predominantly in superficial layers 2/3, but also L6. Sound-evoked activity was decomposed into an early auditory-related component spanning the layers, but most prominently in L2/3 and L5/6, with motor-related activity again mostly contributing in superficial layers. Note how motor activity explained a similar pattern during auditory and visual trials, but shifted to an earlier time window in auditory as compared to visual trials. These results were corroborated by analyses of the local field potential, showing similar onset latencies of auditory and visual evoked activity and predominance of motor-related activity in superficial layers (Ext. Data Fig. 7). When estimating the laminar organization of visual and auditory components with a model-free approach and plotting the onset latency of visual- and auditory-evoked single-unit activity as a function of cortical depth, we found similar results as for the GLM analysis of firing patterns (Ext. Data Fig. 7h). Together, these results indicate different spatiotemporal profiles for visual, auditory, and motor-related components in V1. Our data are most consistent with auditory stimuli evoking fast auditory-related inputs predominantly to deep layers and later, motor-related activity mostly in superficial layers.

### Auditory cortex as a source of early sound-evoked activity in V1

The short-onset, sound-evoked activity in V1 (27 ms), shortly after AC (19 ms, Fig. 3b), suggests that AC may be a source of early auditory-related activity in V1. To test this, we first bilaterally injected AAV-CamKIIa-ChR2-eYFP in AC of naive mice (Ext. Data Fig. 8a). Axonal terminals were observed in layers L1 and L5/6 of V1 (Ext. Data Fig. 8b,c). This AC-V1 projection pattern matches that found in earlier studies (Ibrahim et al., 2016; Rockland and Ojima, 2003). In a subset of animals, cell bodies were photostimulated in AC and the LFP response was recorded in V1 (Ext. Data Fig. 8f-j). Ten millisecond laser pulses over AC led to short-latency LFP deflections in V1, indicative of AC to V1 connections (Ext. Data Fig. 8g), in line with (Ibrahim et al., 2016; Iurilli et al., 2012).

Next, we bilaterally injected AC with muscimol in MST mice during task performance (Fig. 4a). Muscimol infusion immediately abolished spontaneous multi-unit activity in AC (Fig. 4b) but did not affect V1 firing rate as compared to the saline control (Fig. 4c). Muscimol moderately impaired auditory hit rates at maximal saliency (Amax: F(1,19)=6.41, p=0.020; catch: Athr: F(1,19)=4.26, p=0.053; F(1,19)=1.69, p=0.210; n=11 saline, 11 muscimol sessions; Fig. 4d) without effects on visual hit rates (catch: F(1,19)=3.88, p=0.064; Vthr: F(1,22)=0.09, p=0.773; Vmax: F(1,18)=0.43, p=0.522). Reaction times were unaffected, except for a small reduction in Vmax trials (Vthr: F(1,569)=0.16, p=0.686; Vmax: F(1,748)=6.03, p=0.014; Athr: F(1,487)=0.05, p=0.818; Amax: F(1,705)=0.30, p=0.581; Fig. 4e).

**Figure 4:**
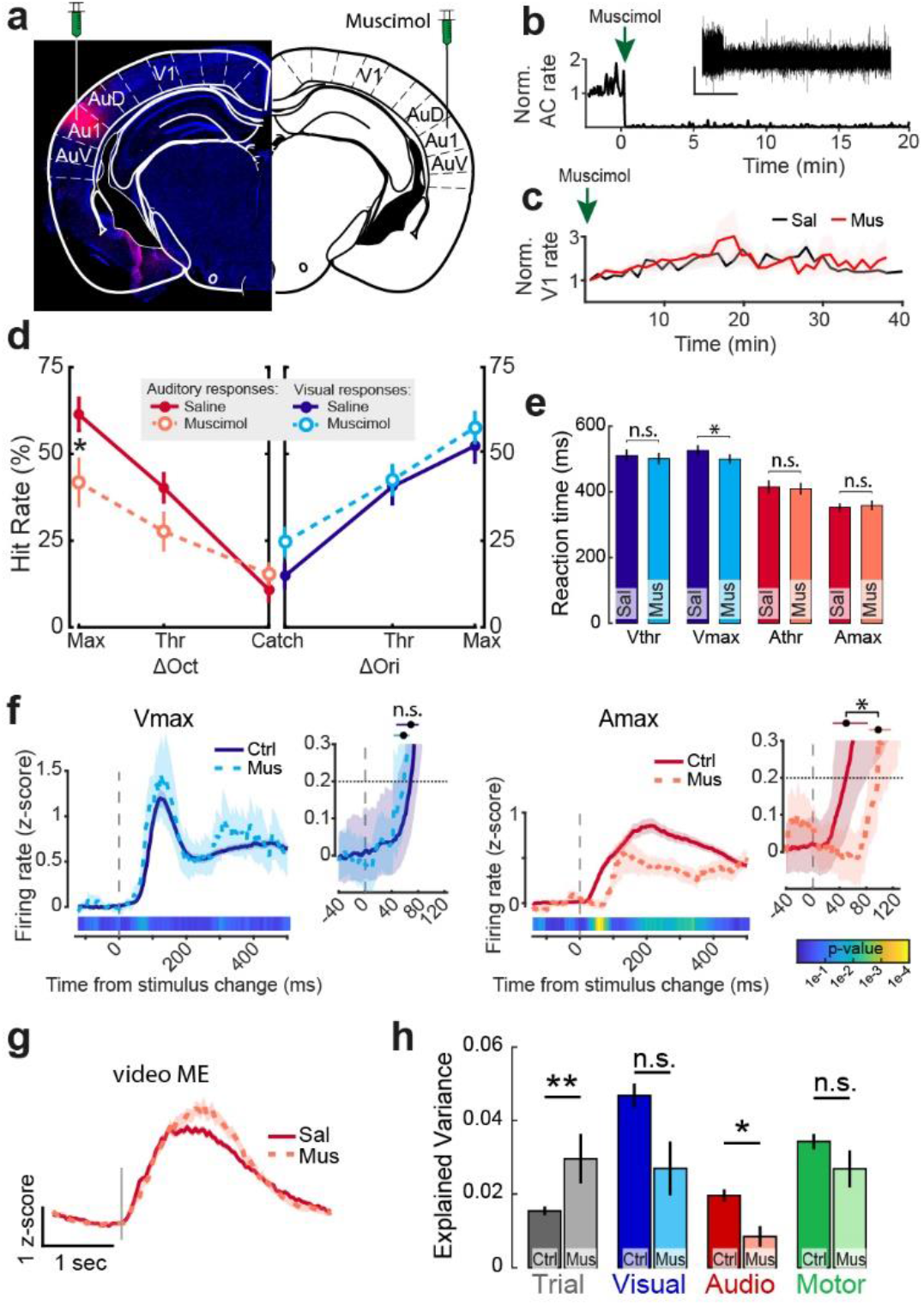
Muscimol in auditory cortex impairs auditory change detection and early evoked activity in visual cortex. **a)** Histological section and schematic of approach to bilaterally inactivate auditory cortex with muscimol. Red blob shows BODIPY TMR-X conjugated muscimol localized mainly in primary auditory cortex (Au1). AuD/AuV: dorsal/ventral secondary auditory cortex. **b)** Multi-unit activity in AC normalized to session start before and after muscimol injection, showing a severe reduction of spiking activity. Inset shows a high-pass voltage trace (cut off at 500 Hz) at one electrode during the same timeframe. Scale bar indicates 100 microvolt and 5 minutes. **c)** Average single-unit activity in V1 normalized to session start for saline and muscimol sessions. Injections were 1-5 minutes prior to recordings. Line and shading indicate mean ± SEM across sessions. **d)** Hit rates in auditory trials (left panel) and visual trials (right panel) for muscimol and saline sessions. Mean ± SEM across sessions. *p<0.05. **e)** Reaction times. Mean ± SEM across trials. *p<0.05. **f)** V1 spiking activity with and without AC inactivation during visual (left) and auditory (right) trials of maximal saliency. Main panels show average spiking activity with the line and shading indicating mean ± SEM across neurons. The lower colormap scales with p-value of bootstrapped test of difference per time bin (n=10.000 bootstraps, p<0.05, Bonferroni corrected; control neurons were subsampled to match the number of recorded V1 neurons during AC muscimol infusion; dataset: 32 control sessions, 570 V1 neurons; 4 muscimol sessions, 53 V1 neurons). The insets show a close up of the early post-stimulus time window and bootstrapped activity (line and shading indicate mean ± 95% confidence interval). The dashed line indicates statistical threshold of deviations from baseline. Upper horizontal dot and error bar indicate when bootstrapped activity significantly exceeded baseline activity. The non-overlap of these distributions showed a significant difference in onset latency (p<0.05). **g)** Overall orofacial motion energy during auditory trials. Grey line: stimulus change. Mean ± SEM across trials. **h)** Explained variance of V1 spiking activity by each predictor subset in the regression model with and without AC inactivation. Mean ± SEM across neurons. *p<0.05. **p<0.01.

Recordings during task performance revealed that the firing rate of V1 neurons following visual stimuli during AC inactivation was similar to control experiments (Fig. 4f). However, AC inactivation affected V1 firing during auditory trials particularly in the first 100 ms and was still associated with a later increase in average firing rate. The onset latency of increased firing was not significantly different for visual stimuli (control: 69 ms (47 - 81 ms) versus muscimol: 58 ms (43 - 67) ms; median and bootstrapped 95% CI; maximal saliency), but delayed for auditory stimuli (control: 50 ms (30 - 82) ms versus muscimol: 98 ms (84 - 117) ms; bootstrap test; p<0.05). Thus, muscimol suppressed the early component of the V1 response to auditory stimuli (from about 30 to 80 ms; Fig. 4f). Muscimol did not affect activity during Athr trials, consistent with the fact that V1 activity during these trials was mostly motor-related, with only a minimal auditory-related component (see Fig. 3f). Given that auditory performance was reduced but not abolished by muscimol injection, the residual auditory hits were still associated with instrumental licking and orofacial movements (Fig. 4g), which could underlie late V1 firing during auditory trials. We tested whether muscimol selectively affected the contribution of auditory predictors to V1 firing rate, while preserving other components (Fig. 4h). Auditory predictors explained significantly less variance under muscimol (p=0.04; Wilcoxon rank sum test), while visual and motor predictors were not associated with significant reductions (Vis: p=0.23; Motor: p=0.95). The variance explained by trial number increased (p=0.01). In sum, AC inactivation selectively affected the early auditory-related component of sound-evoked V1 activity, but not the motor-related component.

### Behavioral dominance of audition over vision

How do auditory and motor signals impact concurrent visual processing? In a subset of sessions in MST mice, we interleaved trials in which both visual and auditory stimuli changed simultaneously with standard trials presenting a unisensory change and catch trials. In these mice, visual and auditory feature changes were associated with different motor actions. The modalities therefore acted as competing inputs and a simultaneous change presented the animal with a conflicting situation. Therefore we first describe the behavior.

In behavioral sessions we presented four levels of visual and auditory feature change that matched in subjective saliency across modalities (Meijer et al., 2018; Song et al., 2017). Stimuli were taken as the x-axis positions corresponding to the same positions along the psychometric function of each modality in unimodal trials and thus matched performance (Fig. 5a; subthreshold (*sub*), threshold (*thr*), suprathreshold (*sup*), maximal (*max*)). Saliency-matched conditions led to comparable increases in pupil dilation (Ext. Data Fig. 9a,b). In conflict trials, animals predominantly chose the auditory lick spout (Ext. Data Fig. 9c). Behavioral choice scaled with the saliency of the sensory input, but in saliency-matched conditions auditory choices dominated (Fig. 5b,c; Ext. Data Fig. 9d). A saliency-matched dominance index (smDI) was computed as a ratio of auditory to visual choices for saliency-matched conflict conditions (Fig. 5d). The smDI was significantly higher than zero (indicating auditory dominance), confirming auditory dominance (smDI: 0.33, Wilcoxon signed rank test, p=0.0014, n=17 mice, based on 97 sessions, 51.932 trials). Audition thus dominates behaviorally over vision in our modality identification task and this is partly explained by faster auditory processing (Ext. Data Fig 9).

**Figure 5:**
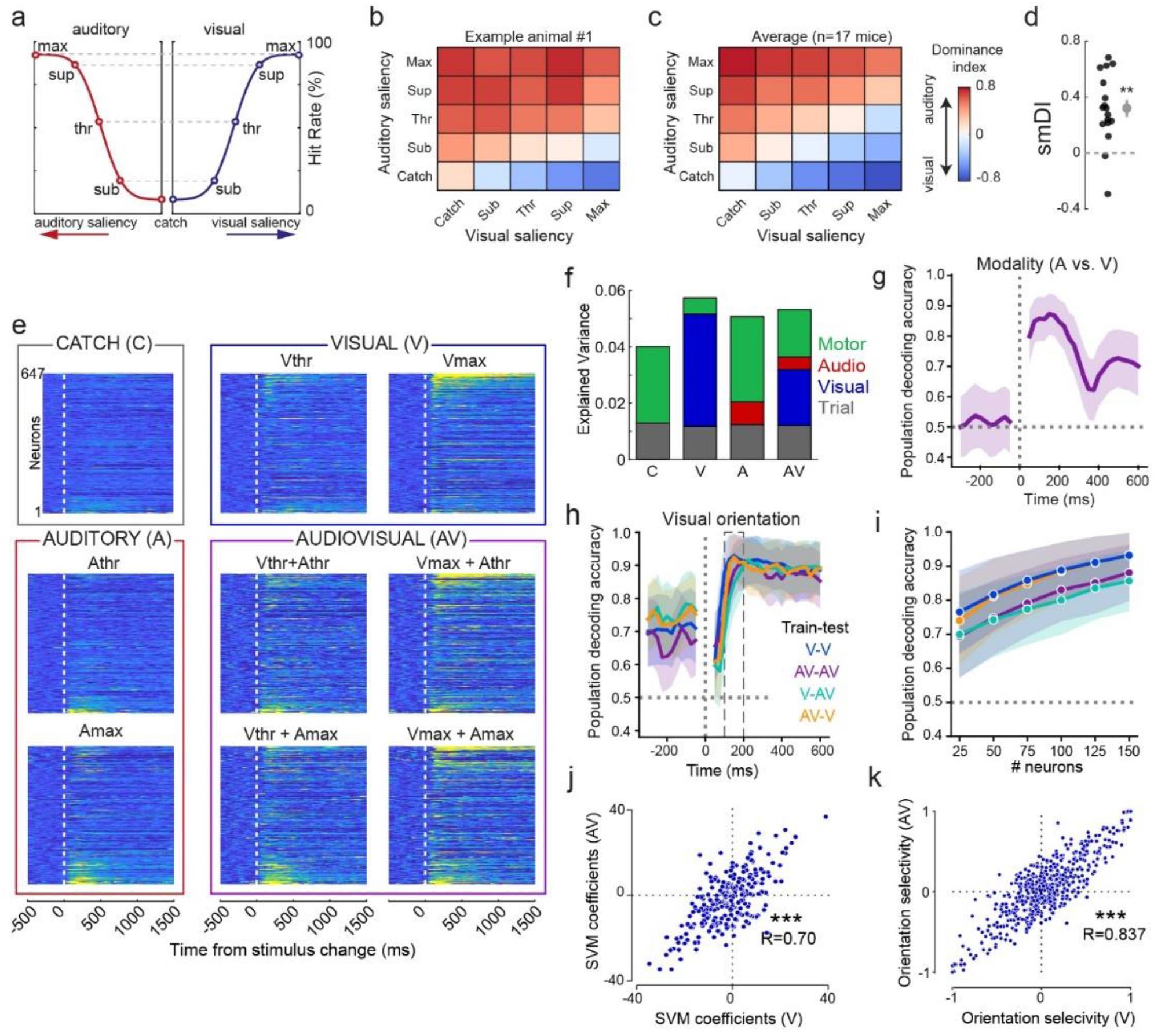
Preserved orientation coding in V1 during multisensory trials showing auditory behavioral dominance. **a)** To equalize subjective saliency during conflict trials, four levels of auditory and visual change were taken at equal positions along this schematic psychometric curve. All data in this figure are from MST animals. **b)** Dominance index (DI) as a heatmap across unimodal and conflict trials for an example animal showing auditory dominance. A positive (red) DI indicates a higher fraction of auditory choices relative to visual. **c)** Heatmap as in (b), but averaged across all animals (n=17). **d)** Saliency-matched Dominance Index (smDI) taken by averaging the DI of all saliency-matched conflict conditions (conditions along top-right to bottom-left diagonal in heatmaps of (b) and (c), excluding catch trials). Each dot is one mouse. Grey dot and error bar are mean ± SEM. **p<0.01. **e)** Heatmap of z-scored firing rate across V1 neurons (each row in each heatmap is a neuron). Each heatmap shows a trial type, with columns increasing in auditory saliency (no change, threshold, maximal) and rows increasing in visual saliency. Lower right 4 panels indicate conflict trials. Neurons are sorted by response magnitude difference between unimodal visual and auditory trials. During audiovisual trials, subsets of visually responsive (upper rows in each heatmap) and auditory responsive neurons (bottom rows) respond similarly as during unimodal trials. **f)** Total explained variance in firing rate of V1 cells for each trial type, with each color indicating the contribution of each predictor subset. Result obtained from the regression model. C: catch trials. **g)** Decoding performance over time for decoders trained to discriminate the modality of unimodal trials (visual versus auditory) from V1 pseudopopulation activity (n=150 neurons). Line and shading indicate mean and 95% CI. Grey dotted line is chance level. **h)** Same as (g), but for performance of decoders trained to discriminate visual orientation. Decoders were trained and tested on held-out test trials of the same trial type (within condition: V-V and AV-AV) or trained on visual and tested on held-out audiovisual trials, or vice versa (cross-condition: V-AV and AV-V). Baseline performance results from a subset of neurons persistently coding orientation, see Fig. 2h,m. Line and shading indicate mean and 95% CI. **i)** Quantification of orientation decoding performance averaged over dashed time window in (h) for different pseudopopulation sizes. **j)** Orientation decoding weights (SVM coefficients) at t=0.175 sec from decoders trained separately on visual or audiovisual trials. Each dot is a neuron. **k)** Orientation selectivity during visual (Vmax) and audiovisual trials (Vmax+Athr or Vmax+Amax).

### Dissociation of auditory and visual processing: preserved orientation coding during conflicting multisensory inputs

To examine how auditory- and motor-related inputs intersect with concurrent visual processing, we sampled neural activity during conflict trials of threshold and maximal audiovisual saliency in 65 out of 122 recording sessions with MST mice. The time window of analysis was broadened to 0-500 ms to investigate continued interaction between visual, auditory and motor-related processing.

We first compared trial-averaged activity during multisensory conflicts with saliency-matched unimodal trial types (Fig. 5e). Neurons that responded during visual or auditory trials continued doing so during conflict trials. In the regression model, visual, auditory and motor-related predictors continued to predict V1 firing rate in held-out conflict trials (Fig. 5f).

These results are in line with our finding that visual, auditory and motor components evoked activity in largely distinct neuronal subsets of V1 (Fig. 3f). Briefly revisiting the unisensory trials, population activity would be expected to distinguish between visual and auditory trials. Indeed a decoder trained to discriminate the modality of unimodal trials from V1 pseudopopulation spiking data (pseudopopulation size n=150 neurons) did so with high accuracy (Fig. 5g).

Returning to conflict trials, the auditory stimulus and orofacial movements explained more variability in conflict trials than visual stimuli did (Fig. 5f). With these multimodal inputs competing, we wondered how visual representations would be affected. We trained the same population decoder to discriminate visual orientation in unimodal visual and multimodal (conflict) trials and tested performance on held-out test data of the same or different trial type (i.e. with or without auditory stimuli). For all conditions tested, orientation decoding performance increased after visual stimulus change (Fig. 5h) and was comparable between visual-only and conflict trials, also when tested for various population sizes (Fig. 5i). The contributions of individual neurons (SVM coefficients) were strongly correlated between visual and audiovisual trials (Fig. 5j; R=0.70, F(1,247)=236.24, p=7.28*10^−38^). This population-level finding matched with single-neuron analyses, where orientation selectivity during visual and conflict trials was strongly correlated in a similar, but even stronger manner (Fig. 5k; R=0.837; F(1,529)=282.61, p=3.97*10^−51^). Therefore, while sounds dominate behavioral choice and V1 variance, visual feature coding is only minimally affected.

## Discussion

Several studies have reported sound-evoked activity in V1, but often without controlling carefully and systematically for sound-evoked behavioral changes. We found that a large part of sound-evoked activity in V1 appeared correlated to orofacial movements, in line with a recent report (Bimbard et al., 2021). However, next to motor-related activity, distinct early auditory-related activity was observed in V1. These inputs likely reflected auditory sensory-evoked inputs as they had short-onset latencies in spiking and LFP data, did not correlate to orofacial movements, and were reduced after AC inactivation (Fig. 2d,e; 3b,f,g; 4f,h; Ext. Data Fig. 7g,h). Rather than there being a single external input of a purely sensory or purely behavioral origin, both auditory and motor-related inputs reach primary visual cortex, and these are segregated in time and space (via different V1 subsets). Sounds can thus lead to strong activity changes in visual cortex through multiple pathways. In addition to the dissociability of auditory and motor processing, we show that auditory and visual streams remain largely segregated, most poignantly illustrated by the preserved orientation coding during multisensory conflict trials in which auditory inputs dominated behaviorally.

### Early auditory-related activity in visual cortex

Even when motor-related influences were accounted for, short-latency AC-dependent signals were present in V1. Onset latencies of sound-evoked activity were comparable to the prior literature with 18 ms in AC (11 ms in Sakata and Harris, 2009) and 27 ms in V1 (35.8 ms in Iurilli et al., 2012). This latency difference is compatible with mono- or disynaptic AC-V1 connections. The specific reduction of auditory-related activity in V1 after bilateral pharmacological inactivation of AC further establishes the efficacy of the AC-V1 pathway during task performance (Fig. 4). Muscimol injections also increased the variance explained by trial number; it may be speculated that this effect was caused by slow fluctuations in excitability of thalamocortical networks affecting V1 activity across the session, even though the mean firing rate was comparable to control recordings (Fig. 4c). AC inactivation impaired, but did not abolish the detection of frequency changes, in line with a modest role for AC in pitch discrimination (Ceballo et al., 2019). Given that primary auditory and visual cortices are monosynaptically connected and sound-evoked activity in V1 has been reported under anesthetized conditions (Henschke et al., 2015; Ibrahim et al., 2016; Iurilli et al., 2012), our findings underscore the efficacy of direct auditory inputs to visual cortex in awake behaving mice, which is apparently not overshadowed by motor and arousal effects or task-dependent factors that could in principle regulate the strength of these inputs (cf. Knopfel et al. 2019). Our anatomical tracing and LFP results are in line with previous reports that sounds can modulate visual cortex through both superficial L1 projections (Ibrahim et al., 2016) and deeper inputs (Iurilli et al., 2012). These inputs may modulate cortical activity through translaminar dendrites (e.g. targeting L1 apical dendrites of L2/3 and L5 neurons) as well as through translaminar inhibitory circuits, modulating activity in supragranular and infragranular layers (Fig. 3) (Ibrahim et al., 2016; Iurilli et al., 2012; Knöpfel et al., 2019; Meijer et al., 2017).

Visually evoked activity in auditory cortex, on the other hand, was strikingly absent. This asymmetry – or predominance of auditory crossmodal influences on visual cortex relative to visual-to-auditory cortex influences - matches earlier physiologial and anatomic studies (Budinger and Scheich, 2009; Campi et al., 2010; Ibrahim et al., 2016; Iurilli et al., 2012; Oh et al., 2014), but is noteworthy because it now is shown to hold in a task setting where auditory and visual changes were of equal behavioral relevance. This asymmetry, however, might also depend on the stimulus characteristics (Chou et al., 2020).

### Behavioral component of sound-evoked activity in visual cortex

A large part of sound-evoked activity in V1 correlated to orofacial movements. Already 60-100 ms after auditory changes, motor activity started (Fig. 2a) which correlated with V1 spiking activity (Fig. 2e; 3f; 5f) and LFP (Ext. Data Fig. 3g). These orofacial movements were themselves stimulus-specific, such that auditory stimulus identity could be decoded from video footage (Fig. 2j-l). Although the auditory stimuli were a weighted combination of different tones, ‘motor-tuning’ might arise through differences in subjective saliency, arousal, or aversion to particular auditory frequencies. These results, including the frequency-tuned responses in V1, are in agreement with a recent study showing that different sound clips were associated with stereotypical orofacial movements across mice, which correlated to V1 activity (Bimbard et al., 2021). These and the present results strongly argue for cautious interpretations of multisensory interactions in awake subjects and underline the need to carefully monitor behavioral state.

We labeled neural activity changes related to orofacial movements as motor-related. It is unclear whether these are better interpreted as corollary discharge signals to V1 predicting (or otherwise relating to) visual consequences of motor movements (Guitchounts et al., 2020; Leinweber et al., 2017; Pennartz et al., 2019; Schneider et al., 2014), or as internal state changes associated with arousal levels, which may be linked with increased movement (Niell and Stryker, 2010; Vinck et al., 2015). Visual and auditory stimuli evoked approximately similar levels of pupil dilation (Ext. Data Fig. 9a), but auditory stimuli could in addition elicit fast arousal responses, known to originate from intralaminar thalamic nuclei (Minamimoto and Kimura, 2002; Van der Werf et al., 2002). As corollary discharge signals might be temporally shifted relative to video-observed movements (Leinweber et al., 2017), a methodological limitation is that our regression model does not take into account nonlinear or temporally shifted relationships between orofacial movements and V1 activity. Whether sounds predominantly activate arousal or sensorimotor signalling in visual cortex is a question for future investigation.

### Preserved orientation coding during auditory behavioral dominance

Mice reported auditory over visual stimulus changes during saliency-matched conflict trials in a similar manner as in Song *et al*. (2017) (but see (Coen et al., 2021) for balanced audiovisual weighting). During these conflict trials, the combination of auditory and motor-related components explained more variability in firing rate than visual stimuli, even though orientation coding was preserved. In other words, both auditory and motor influences on V1 activity appear to be largely orthogonal to visual feature representations, cf. (Montijn et al., 2016; Stringer et al., 2019). This preserved orientation coding matches well with our observation that visual, auditory, and motor-related activity occurs in rather segregated cell populations (Fig. 3f) and underscores their dissociability. Primary visual cortex thus supports (relatively independent) parallel encoding of signals related to different sensory and motor modalities. This segregation somewhat contrasts with studies reporting predominantly jointly responsive neurons (Bizley et al., 2007; Knöpfel et al., 2019), which may relate to our behavioral task in which auditory and visual cues were explicitly not to be integrated.

### Beyond cue integration

Our work differs in an important way from the large body of literature on multisensory cue integration (Fetsch et al., 2013; Meijer et al., 2019; Stein and Stanford, 2008), where for instance crossmodal inputs are interpreted to improve the inference of grating orientation (Ibrahim et al., 2016; Nikbakht et al., 2018; Williams et al., 2021). In the MST task, the two modalities were not jointly informative about the same external variable (e.g. they were not jointly indicating a source location, heading direction or stimulus rate), but rather auditory signals were statistically uninformative about visual features (and vice versa), in contrast to e.g. (Garner and Keller, 2022; Lippert et al., 2007; Meijer et al., 2020, 2018; Sheppard et al., 2013). Instead, animals needed to monitor potential changes in both modalities across time, without a trial onset cue, and discriminate in which modality a change occurred.

Our finding that auditory changes not only modulated, but evoked spiking in V1 of naive and visually trained mice as well as in mice trained to discriminate auditory and visual signals, suggests a broader role for auditory signals in V1 than only sharpening visual tuning. As shown elsewhere, auditory signals in V1 do not become causally important for audition during MST training, as V1 optogenetic inhibition impacts visual but not auditory change detection performance (Oude Lohuis et al., 2022). Even if auditory inputs to visual cortex are not directly relevant for detecting single visual features, they are hypothesized to fit in a broader view of sensory cortical function (Meijer et al., 2019; Pennartz, 2015, 2009; Petro et al., 2017), where crossmodal interactions serve to orchestrate perception across a larger cortical network, guide crossmodal attention, and inform visual processing in distributed networks about ongoing auditory events. In this respect, it is interesting that the auditory component was stable across cohorts (Ext. Data fig. 5c), consistent with a basal, rather than task-specific function. Part of this orchestration may reside in predictions that fast auditory processing conveys upon vision; another part may relate to the observation that perceptual phenomenology is qualitatively rich, and thus requires segregation as well as integration of sensory modalities (Pennartz, 2015, 2009).

In sum, to correctly interpret (multi)sensory-evoked activity, careful dissociation of sensory and motor origins is necessary. Through this dissociation it becomes clear that sound evokes inputs from auditory to visual cortex that are fast and transient, as well as to later, secondary motor modulations through sound-evoked body movements. The associated activity patterns temporally overlap somewhat and co-exist with visual processing, although this remains dissociable as apparent from population coding of visual grating orientation. An exciting direction of future research will be to understand how these multiple signals co-exist to contextualize sensory input to generate meaningful information processing and behavior.

## Acknowledgments

We thank Jorrit Montijn and Guido Meijer for useful comments on the manuscript, D. Sridharan for providing code for the multi-alternative detection model; C. Rossant, members of the Cortex Lab (UCL) and contributors for Klusta and Phy spike sorting software; Andriana Mantzafou, Klara Gawor, and Alexis Cervàn Canton for assistance in behavioral training. This work was supported by the European Union’s Horizon 2020 Framework Program for Research and Innovation under the Specific Grant Agreement 945539 (Human Brain Project SGA3) to C.M.A.P. and by the FLAG-ERA JTC 2019 project DOMINO (co-financed by NWO) to U.O.

## Author contributions

Conceptualization, M.O.L., U.O., C.M.A.P.; Methodology, Formal Analysis, Visualization, M.O.L.; Additional analysis, P.M.; Writing, M.O.L., U.O., C.M.A.P.; Supervision & Funding, U.O., C.M.A.P.

## Data availability

All behavioral and neural data related to this study will become available on FigShare upon publication.

## Code availability

All code related to this study become available on GitHub upon publication.

## Methods

### Animals

All animal experiments were approved by the Dutch Commission for Animal Experiments and by the Animal Welfare Body of the University of Amsterdam. Thirty-three male mice were used from different genotypes: wildtype C57BL/6, PVcre (JAX 008069), and PVcre/TdTomato (JAX 027395). Mice were at least 8 weeks of age at the start of experiments. Mice were group-housed under a reversed day-night schedule and all experimental procedures were performed during the dark period (8:00 – 20:00).

### Head bar implantation

Before the start of any experiment, a custom-made titanium head-bar was implanted to allow head fixation. Mice were anesthetized with isoflurane and fixed in a stereotaxic apparatus. A circular patch of skin was removed to expose and disinfect the skull. A circular head bar (inner diameter 10 mm) was positioned over the skull to include bilateral V1 and AC and glued and cemented to the exposed skull. After a recovery period of 2-7 days, mice were habituated to handling and head-fixation before start of the training procedure.

### BEHAVORIAL TASKS

Mice were water-deprived throughout the course of experiments and earned their daily ration of liquid by performing the behavioral task. In the case of low performance, daily intake was supplemented to a minimum of 0.025 ml per gram of body weight. During a session, mice were headfixed their bodies were positioned in a cylindrical holder. Two lick spouts were positioned symmetrically on the left and right side within reach of their tongue. Licks were detected by capacitance-based (during training) or piezo-electric based detectors (during recordings). Correct licks were immediately rewarded with 5-8 μl of liquid reward (infant formula; Nutrilon) delivered through the same lick spout using gravitational force and solenoid pinch valves (Biochem Fluidics, Boonton, USA).

### Audiovisual change detection task

In the audiovisual change detection task, auditory and visual stimuli were continuously, without pre-cueing, presented throughout a behavioral session. During visual trials a feature (the orientation) changed, after which this new feature (the post-change orientation) continued to be shown. Similarly, auditory trials consisted of a change of one auditory stimulus to another.

### Stimuli

Visual stimuli were drifting square-wave gratings with a temporal frequency of 1.5 Hz and spatial frequency of 0.08 cpd at 70% contrast (35 cd/m^2^ luminance difference between bright and dark). In trials with a visual change the orientation of the drifting grating was instantaneously changed (e.g. from 60° to 90°) while preserving its phase. Visual stimuli were presented with a 60 Hz refresh rate on an 18.5-inch monitor positioned at a straight angle with the body axis from the mouse at 21 cm from the eyes.

Each auditory stimulus was a weighted combination of five pure tones at harmonic frequencies: a center tone, as well as two lower and two higher harmonics (octaves below and above the center tone). If f_0_ is the center tone, then: *f*_*-2*_ = ¼\**f*_*0*_, *f*_*-1*_ = ½\**f*_*0*_, *f*_*0 =*_ *f*_*0*_; *f*_*+1*_ = 2\**f*_*0*_; *f*_*+2*_ = 4\**f*_*0*_. We name each auditory stimulus after the frequency of its center tone. All frequencies were expressed in scientific pitch as powers of 2 with the center tones spanning from 2_13_ Hz (=8372 Hz) to 2^14^ Hz (=16744 Hz). An example stimulus, 2^13.5^ (named by center tone), was therefore composed of five pure tones of 2^11.5^, 2^12.5^, 2^13.5^, 2^14.5^, and 2^15.5^ Hz. The weight with which each tone was present was taken from a Gaussian distribution across all tones for all stimuli, centered at 2^13.5^ (=11585 Hz). Lower and higher harmonics thus contributed less to the auditory stimulus than the center tone. Because of this fixed weight distribution, stimuli with higher center tone frequency have decreasing weights for higher harmonics and increasing weights for lower harmonics. Stimuli with higher center frequency are thus increasingly made up of lower frequency components to the point of arriving at the starting stimulus (see also Ext. Data Fig. 1). This auditory stimulus design with harmonics and fixed weights was inspired by the Shepard tone illusion (Shepard, 1964). However, in contrast to this illusion, our stimuli were static and not sweeping across frequencies, and the original illusory aspect of a tone ever-increasing (or decreasing) in pitch was not exploited. The primary reason for this auditory stimulus design was the circular nature of the stimulus set, which mirrored the visual stimulus set with drifting gratings in all orientations.

During trials with an auditory change, one stimulus was changed instantaneously to another. This resulted in a shift in spectral power to five new frequencies which appeared to the mouse as an increase or decrease in pitch. Auditory changes were expressed as partial octaves, with ½ octave maximally salient and the minimal change used was 1/256 partial octave. The degree of frequency/octave change determined the auditory saliency and was varied across experimental conditions. During auditory stimulus changes, the phase across all tones was preserved. Stimuli were presented with a sampling rate of 192 kHz. Stimuli were high-pass filtered (Beyma F100, Crossover Frequency 5-7 kHz; Beyma, Valencia, Spain) and delivered through two bullet tweeters (300 Watt) directly below the screen. Note that this high-pass filter eliminated the lowest frequency components of the Shepard stimuli, and left the mid and high frequency components intact (those that span the sensitive part of the mouse hearing range, 8-16 kHz). This was done to prevent damage to the specialized tweeters that we used, but did not affect the animals’ ability to report even very small differences between subsequently presented Shepard tones. Sound pressure level was calibrated at the position of the mouse and volume was adjusted per mouse to the minimum volume that maximized performance (average ±70 dB).

In an earlier cohort of mice (N=13/33) and for the audiovisual detection task (n=3, see below), the Shepard tones (1) were expressed in absolute Hz (e.g. an auditory trial with Δ2kHz changed from 8 kHz to 10 kHz), (2) had 9 instead of 5 harmonics, (3) were presented with a sampling rate of 48 kHz and (4) were not phase-preserved during a change in auditory frequency. We observed no qualitative or quantitative differences in both neural and behavioral results between the cohorts and the data was pooled for all analyses.

### Trial types

Trials simply consisted of an instantaneous feature change (visual, auditory or audiovisual) and an ensuing reward window. Trial onset was defined by an instantaneous change in the visual or auditory stimulus, or no change (catch trial). All analyses are relative to this stimulus change. Trials were separated by an inter-trial interval randomly taken from an exponential distribution (mean 6, minimum 3, and maximum 20 seconds). Trial types were pseudorandomly ordered by block-shuffling per 10 trials (8% catch trials=no change, 46% visual trials, 46% auditory trials). In sessions with multimodal conflict trials these replaced unimodal trials (see below).

### Task versions

Animals were assigned to one of three versions of the audiovisual change detection task in which the visual and auditory stimuli were identical and only the reward contingencies varied (i.e. which stimuli were rewarded). This led to a controlled manipulation of the behavioral relevance of the stimuli as well as differences in the amount of instructed movements (goal-directed licking). Noncontingently exposed (NE, n=7) animals were not rewarded for licking after auditory or visual stimulus changes, but obtained rewards for licks during hidden ‘response windows’ that were temporally offset from the stimuli. This resulted in spontaneous licking behavior at the two spouts that was occasionally rewarded. Unisensory-trained (UST, n=4) animals were trained to only report visual changes and ignore auditory changes. Auditory stimuli and changes were presented throughout all sessions, but were not associated with reward and changes were temporally decorrelated from the task-relevant visual trials (no accidental conflict trials were programmed). Multisensory-trained animals (MST, n=17) animals were trained to detect and identify one of both modalities. Animals were required to respond in a lateralized manner to each modality: lick to one side to report visual changes, to the other side for auditory changes (modality-side pairing was counterbalanced across mice). In other words, mice were required to simultaneously monitor both the auditory and visual modality and identify the sensory modality in which a change occurred. As we performed additional experiments with animals from the MST cohort, this resulted in a higher number of animals in the MST cohort.

### Psychometric performance

Animals in the NE cohort were accustomed to spontaneous licking behavior irrespective of sensory stimuli in a few sessions. Animals in the UST and MST cohorts were trained over the course of several weeks in which progressively more difficult trial types (lower saliency) were introduced and reward size was lowered until performance stabilized. To match the subjective salience of auditory and visual stimuli across mice and modalities we chose intensities according to their unimodal hit rates (Meijer et al., 2018; Song et al., 2017). For each trained animal we established perceptual sensitivity by presenting five levels of auditory and visual saliency (amount of change) that spanned the perceptual range for three consecutive behavioral sessions. We fit the concatenated data of these three sessions with a cumulative normal distribution per modality with four free parameters (Meijer et al., 2018):

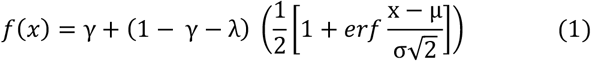

Here, γ describes the false alarm rate (spontaneous licks during catch trials), λ the lapse rate (misses at maximal saliency), μ the mean of the cumulative normal distribution (perceptual threshold), and σ the standard deviation (sensitivity to variations of stimulus intensity). Having established the psychometric function per mouse, we took four levels of saliency per modality at fixed points along the psychometric function: subthreshold (μ-σ; *sub*), threshold (μ; *thr*), suprathreshold (μ+ σ; *sup*), and maximal saliency (*max*).

### Conflict trials

In a subset of sessions we introduced multimodal trials in which auditory and visual stimuli simultaneously changed (conflict trials, making up 25% of all trials; replacing a fraction of unimodal trials). These multimodal trials were introduced in experiments with NE and MST mice. Multimodal trials were omitted in experiments with UST mice as systematic pairing of auditory and visual changes would render the auditory stimuli predictive of concurrent visual changes, while in these mice we aimed to study the processing of auditory stimuli under behaviorally irrelevant conditions.

For NE mice, multimodal trials were presented during nearly all recording sessions (n=27/28 sessions). For MST mice, we first quantified behavioral choice during conflict trials in a set of sessions without recordings for all combinations of visual and auditory saliencies (4×4=16 saliency combinations). This protocol was repeated for 4-7 sessions per animal and the data was averaged for analysis (total N=97 sessions; N=17 mice). For MST mice, the side of the first lick was registered as the animal’s choice. To maintain consistency in task rules (i.e., that a change predicts reward) licking to both spouts was rewarded. In a separate set of experiments (N=20 sessions, 4 mice) we systematically varied the stimulus-onset-asynchrony (SOA) of the auditory and visual change in conflict trials. In these trials we presented only one combination of auditory and visual subjective intensity (both at *threshold* saliency) and used the temporal offsets: -300 ms, -100 ms, -30 ms, 0, +30 ms, +100 ms, +300 ms (where negative values mean that the auditory stimulus changes first). Multimodal trials were also introduced in a subset of the recording sessions (N=65/122 recording sessions). During recording sessions only two levels of saliency were used.

Due to our continuous stimulus design, we were constrained in the timing of visual and auditory changes. A constraint on the auditory change resulted from the fact that the auditory stimulus was changed only when all component tones were aligned in phase. This was done to avoid inducing artefacts changing the frequency of pure tones out of phase. For the visual domain, a constraint on precise timing resulted from the refresh rate of the monitor (60 Hz). To achieve maximal alignment in audiovisual trials, we first computed the future timestamp of the phase-preserved change in auditory frequency. The visual stimulus changed at the frame closest to that timestamp. Stimulus-onset asynchrony was therefore maximally 8.33 ms; half the duration of the interframe interval (0.5 * 16.67 ms). The direction of this misalignment varied and was small relative to the timescale of analysis of conflict trials (0-500 ms).

### Stimuli during recording sessions

During recording sessions, the trial type conditions were limited to get sufficient repetitions. First we limited trials to use only two levels of saliency, *threshold* and *maximum*. Threshold intensity was obtained through psychophysical experiments per modality and per mouse (described earlier) and maximum intensity was always fixed (90 deg and ½ octave). Second, we fixed the auditory and visual stimulus identities, such that changes occurred only between 4 visual stimulus orientations and 4 auditory frequencies (A, B, C, D). The distance between A and B and between C and D was at threshold level, while the distance between A and C and between B and D was maximal. An example stimulus set was 90, 97, 180, 187 (in degrees), and 2^13^, 2^13.03125^, 2^13.5^, 2^13.53125^ (in partial octaves; Ext. Data Fig. 1d,e). Auditory and visual stimuli therefore jumped back and forth between four orientations and frequencies across trials, providing reliable estimates of tuning to specific features. For naive mice we used threshold values that matched those from trained animals. Nearby stimuli (A and B, as well as C and D) evoked highly similar activity patterns and for all analyses, the two nearby stimuli were grouped (i.e. set A/B and set C/D).

### Engaged versus passive

In a subset of MST mice and in a separate set of sessions (N=5), we combined task-engaged and passive blocks within the same recording session. During active blocks the lick spouts were accessible, while in passive blocks the lick spouts were manually positioned out of reach. The stimuli as well as the temporal statistics and trial type distributions were the same for the active and passive blocks within that recording session. In some sessions we implemented one passive and one active block. In other sessions multiple active and passive blocks were alternated (n=5-7 blocks). The order of passive and active blocks was counterbalanced across sessions. No differences were found between single versus multiple alternating blocks and data were pooled. Only video data was analyzed (Ext. Data Fig. 2c-e).

### Audiovisual stimulus detection task

We trained additional mice (n=3) on a simpler variant of the change detection task, in which we could present a larger set of stimuli. In this detection task animals had to detect stimulus presence, rather than stimulus change. Each trial consisted of a blank intertrial interval (gray screen, no sound) drawn randomly from the same distribution as the change detection task and a stimulus window (1.5 seconds) during which a reward could be obtained for licking the correct lick spout. Analogous to the change detection task, animals had to report presence of a stimulus and its modality by directed licks to either the left lick spout (visual) or right (auditory). Modality-side pairing was the same for the three animals. Four trial types were used: visual (41% of trials), auditory (41%), catch (no stimulus, 8%), and conflict trials (10%). Conflict trials were not analyzed. The same stimulus set (full-field drifting gratings and Shepard tones) was used as in the change detection task. The saliency of each trial was now determined by grating contrast, while auditory saliency was determined by sound volume.

Similar to the change detection task, we first established psychophysical performance of each mouse in a series of behavioral sessions with variable visual and auditory saliency across the perceptual range (Meijer et al., 2018). Subsequently, for recording sessions a fixed saliency was chosen at threshold level in the same way as for the change detection task, resulting in substantial numbers of hits and misses. Eight orientations (spaced 45°) and eight frequencies (center tone spacing at 1 kHz) were presented at this saliency level. Visual contrast levels for the three animals were 9.7%, 10%, 13%. Auditory saliency levels were 56, 78, 82 dB. Note the higher volume used for two animals. This cohort of animals was of old age at the time of experiments (40-43 weeks) and two of these animals showed a progressive decline in their sensitivity to auditory stimuli of lower and intermediate volumes. This is in line with age-related hearing loss reported in C57BL/6 mice (Henry and Lepkowski, 1978; Spongr et al., 1997). We included these mice irrespective of their decreased hearing sensitivity as we titrated auditory intensity to result in similar subjective saliency (hit rates were comparable for both visual and auditory trials: 49% and 46%, respectively) and we focused in this task on correlated feature tuning of V1 neurons, not on performance aspects.

### Electrophysiological recordings

On the day before the start of extracellular recording sessions, mice were anesthetized with isoflurane and small (∼200μm) craniotomies were made using a dental drill over the areas of interest. Areas of interest were binocular V1 (relative to lambda: AP 0.0, ML ± 3.00 mm) and AC (relative to bregma: AP -2.6 mm, ML ± 4.3 mm). Craniotomies and recordings in medial prefrontal cortex and posterior parietal cortex were also performed, but data from these areas were not analyzed here. Extracellular recordings were performed on consecutive days with a maximum of 4 days per mouse to minimize damage to the circuitry. Each recording session, up to three microelectrode arrays (silicon probes) of 32 or 64 channels (NeuroNexus, Ann Arbor, USA – A1×32-Poly2-10mm-50s-177, A4×8-5mm-100-200-177, A1×64-Poly2-6mm-23s-160) were slowly inserted into their target area. We approached V1 perpendicularly to the cortical surface and lowered the silicon probe until all recording sites spanned the cortical layers of V1. Because of the circular headbar (inner diameter 10 mm) we used, craniotomies were located slightly medially on the skull surface and AC was approached with an angle approximately 30° away from the midline and with 64-channel laminar probes that spanned 1450 μm. Due to the span and angle of approach, we recorded multiple subfields of the auditory cortex. In a subset of animals, on the last day of recordings the probe was covered in DiI (ThermoFisher Scientific) to facilitate post hoc reconstruction of the electrode tract. Neurophysiological signals were pre-amplified, bandpass filtered (0.1 Hz to 9 kHz), and acquired continuously at 32 kHz with a Digital Lynx 128 channel system (Neuralynx, Bozeman, MT). The start of the behavioral task commenced at least 15 minutes after probe insertion to allow for tissue stabilization.

### Video monitoring

A near-infrared monochrome camera (CV-A50 IR, JAI, Copenhagen, Denmark) was coupled to a zoom lens (50 mm F/2.8 2/3” 10MP, Navitar, Rochester, USA) and positioned at approximately 30 centimeters from the mouse to capture the lick spouts and face of the mouse within a single view. The left side of the face was illuminated with an off-axis infrared light source (IR-LEDs, 850 nm) positioned to yield high contrast illumination of both the eye and whisker pad. A frame grabber acquired images of 752×582 pixels at 25 frames per second. With this acquisition rate the timing precision for the facial motion and pupil size and location was about 40 ms.

### Optogenetics

Mice (N=5 NE animals, 20 weeks old) were subcutaneously injected with the analgesic buprenorphine (0.025 mg/kg) twenty minutes prior to surgery. During surgery, mice were maintained under isoflurane anesthesia (induction at 3%, maintenance at 1.5–2%). We aimed at infecting bilateral AC and centered our injection at primary auditory cortex (A1). We performed a small craniotomy over bilateral primary auditory cortex (A1 relative to bregma: AP -2.60 mm, ML ± 4.30 mm) using an ultra-fine dental drill and inserted a glass pipette backfilled with AAV2-CamkIIa-hChR2(H134R)-EYFP (titer: 3×10¹² viral genomes per ml, 26969-AAV2, Addgene). AC was approached similar to extracellular recordings. Four injections of 13.8 nl were made using a Nanoject pressure injection system (Drummond Scientific Company, USA) at two depths: two at 1200 μm and two at 1000 μm below the dura (ending up in A1 because of the angle). Each injection was spaced apart from the next one by 5 minutes to promote diffusion and prevent backflow. After viral injections, the recording chamber was covered with silicon elastomer (Picodent Twinsil) and mice were allowed to recover.

After 4-6 weeks to allow for viral expression, the silicon elastomer was removed during recording sessions. To photostimulate AC, a fiber-optic cannula (inner diameter 200 um, numerical aperture 0.48, DORIC Lenses, Quebec, Canada) was positioned directly over AC. The fiber-optic cannula was sealed with black tape, leaving only the tip exposed to prevent light from reaching the eye of the mouse. A fiber optic patch cord connected the cannula to a 473 nm laser (DPSS 473nm H300, Eksma Optics, Vilnius, Lithuania). A shutter (LS6 Uniblitz, Vincent Associates, Rochester, USA) controlled light delivery and was located in a sound-insulated box distal from the experimental setup to prevent any auditory-evoked activity. AC was stimulated with 10 ms pulses with variable laser power (0-20 mW total power) and variable frequencies (5, 10, 20 and 50 Hz).

### Muscimol inactivation of AC

On the day before the start of muscimol experiments a craniotomy over AC was made using the same coordinates as for the recordings and optogenetics. To inactivate AC, 300 nl of muscimol solution (10 mM in saline, pH 7.2; Sigma Aldrich) or saline solution (control) was injected in bilateral AC. AC was approached with the same coordinates (centered at primary auditory cortex) and angle of approach as with extracellular recordings or viral injections. Glass micropipettes were backfilled and slowly inserted through the craniotomy. Three injections of 50 nl were done at 1200 μm and three at 1000 μm below the dura using the Nanoject injection system, with one minute spacing between each injection. AC inactivation was verified using multi-unit recordings in A1, taking V1 as a control area. In a subset of animals we injected BODIPY TMR-X conjugated muscimol (ThermoFisher Scientific; Catalog number: M23400) during the last session to assess the localization and spread of AC muscimol injections. Injecting 300 nanoliter led to localized expression in AC, primarily in A1 (Fig. 4a), whereas we found that injecting a larger volume of 500 nanoliter led to reduction of spontaneous activity in V1 after approximately 10 minutes, which we interpreted as extended diffusion.

During experiments with simultaneous recordings the silicon probe was first inserted to stabilize, then we performed muscimol injections, and directly after these the behavioral experiment was started. Muscimol or saline injections were performed on alternating days. We observed comparable behavioral performance on days after muscimol experiments.

### Histology

At the end of the experiment, mice were overdosed with pentobarbital (>100 mg/kg) and perfused (4% paraformaldehyde in phosphate-buffered saline, pH 7.2). The brains were recovered for histology to verify viral expression and recording sites. Coronal sections were cut at 50 μm and overlaid with the matching reference section from the atlas (Paxinos and Franklin, 2004). Flattened cortical sections were cut at 50 μm, prepared as described previously (Lauer et al., 2018), and V1 and AC were identified based on cell densities aligned to reference maps (Gămănut et al., 2018).

### DATA ANALYSIS

Unless otherwise stated, all data were analyzed using custom-made software written in MATLAB (The MathWorks, Natick, USA) or Python (analysis on population decoding only). All code and data will be made available on Github (insert link) and FigShare. Given that sound-evoked activity consisted of both auditory and motor-related activity in V1 of mice from all three cohorts, data from all cohorts were combined, unless otherwise specified.

### Video analysis

To capture and describe orofacial movements from video recordings, the principal components of motion across the video frames were extracted using FaceMap (Stringer et al., 2019). Briefly, the video was spatially downsampled (1 every 4 pixels) and singular value decomposition was applied on the frame-to-frame pixel intensity differences of a representative excerpt of frames (4000 frames). Subsequently, frame-to-frame motion of all frames was projected into the first 500 principal components. Total video motion energy (video ME) was taken as the absolute sum across all 500 components. To investigate the relationship between neural measurements and more detailed orofacial movements, the first *n* principal components were selected that captured movements explaining most of the frame-to-frame pixel intensity differences. For the regression model this was n=25 PCs, and for population feature decoding this was n=30 PCs. The first 25 PCs captured roughly 62% of the variance in frame-to-frame pixel intensity differences.

Pupil size and position were extracted using DeepLabCut (Mathis et al., 2018). The network was trained on 300 frames from 15 video excerpts of 1-2 minutes with varying illumination, contrast, pupil size, imaging angle, and task conditions. We labeled the pupil center and 6 radially symmetric points on the edge of the pupil. An ellipsoid was fit to these 6 outer points. The x and y coordinates of pupil center were taken as the center of the ellipsoid and pupil area as the ellipsoid area from the fitted ellipse parameters. Poorly fitted frames (likelihood <0.9999, output DeepLabCut) were replaced by the running median (median of 10 good frames), except if more than five adjacent frames were poorly fit (e.g. during extended periods of eye closure). We z-scored the total session traces.

### Behavioral dominance

Behavioral dominance was quantified per trial condition by computing a behavioral dominance index (BDI):

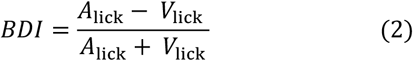

where A_lick_ and V_lick_ are the amount of conflict trials in which the animal chose the auditory or visual lick spout, respectively. Note that misses are not taken into account in this index. BDI values range from +1 to -1, where +1 means exclusively auditory choices and -1 exclusively visual ones. The saliency-matched BDI (smBDI) was obtained by averaging the BDI of saliency-matched conflict trials (A_sub_ + V_sub_, A_thr_ + V_thr_, etc.). To describe and determine behavioral dominance as a function of audiovisual SOA we fitted the behavioral data with a cumulative Gaussian function

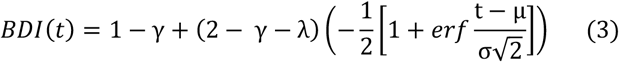

Here, γ is the asymptotic visual dominance, λ the asymptotic auditory dominance, μ the mean of the cumulative Gaussian (time point of crossover), and σ the standard deviation (sensitivity to variations in SOA). We fitted the data using MATLAB’s *fit* function and constrained μ between -300 and +300 ms, σ between 0 and 200 ms, and γ and λ between 0 and 1. Bootstrapped 95% CI was computed from n=1000 fits on resampled data.

### Neural data processing

Before spike sorting the median of the raw trace of nearby channels (within 400 μm) was subtracted to remove common noise artefacts. Automatic and manual spike sorting were done using Klusta and the Phy GUI, respectively (Rossant et al., 2016). During manual curation each putative single unit was inspected based on its waveform, autocorrelation function, and its firing pattern across channels and time. High-quality single units were included as having (1) an isolation distance higher than 10 (Schmitzer-Torbert et al., 2005) (2) less than 0.1% of their spikes within the refractory period of 1.5 ms (Bos et al., 2017; Vinck et al., 2016), (3) stable presence throughout the session. This latter was quantified by binning the firing across the entire session (approximately 45-75 minutes) in 100 time bins and only including neurons that spiked in more than 90 time bins.

To compute firing rates, spikes were binned in 10 ms bins and convolved with a causal half-Gaussian window with 50 ms standard deviation, unless stated otherwise. For analyses where neurons were compared, the firing rate of each single unit was z-scored by subtracting for each trial the mean firing rate of the baseline period (−1 to -0.1 seconds before stimulus change) and dividing by the standard deviation of all baseline periods.

For the initial assessment of how many neurons were significantly modulated after visual or auditory stimulus changes, the firing rate during baseline (−1000 to 0 ms) and post-change window (0-200 ms post stimulus) was compared with a paired two-tailed Wilcoxon signed rank test (p<0.025). Neurons were deemed significantly visually responsive if the firing rate was significantly different for at least one of the two grouped orientations (i.e. A/B or C/D) during maximal saliency trials (and similarly for auditory trials). Only conditions with at least 10 trials were tested. The fraction of significantly responsive neurons was only computed for sessions with at least 15 simultaneously recorded V1 neurons.

The onset latency of spiking activity for individual neurons was estimated using ZETA, a recently developed bin-less statistical test for determining whether a neuron shows a time-locked modulation of spiking activity (Montijn et al., 2021). We opted for this as visual and auditory stimuli can elicit very different neural dynamics in visual and auditory cortex (specifically, spiking responses in A1 can be very brief (DeWeese et al., 2003)) and ZETA prevents confounds related to different temporal dynamics by avoiding the need to bin spikes. ZETA was computed over time for auditory and visual maximal saliency trials. Neurons were deemed significantly modulated if ZETA exceeded a value of 2 during 0-1000 ms after stimulus change. The onset of this spiking response was taken as the onset latency. For Ext. Data Fig. 7g, we focused on sensory-evoked spiking and to minimize occlusion by motor-related confounds, we excluded auditory and visual spiking activity with onsets occurring later than 200 ms (note, however, that this approach does not strictly separate sensory from motor-related activity).

To estimate the onset latency of visually or auditory induced spiking activity with greater temporal detail, the spiking activity was pooled across neurons in an area. Only sessions with at least 10 neurons were included. The spike train of each neuron was divided across 1 ms bins and smoothed with a causal half-Gaussian window with 10 ms standard deviation. The activity was averaged over trials of interest (Vmax or Amax trials). To compare across sessions, the firing rate was averaged across all simultaneously recorded neurons in each area and z-scored, as described for single neurons. The first time bin this z-scored activity crossed a threshold of 1 was taken as the onset latency of the population activity.

### Feature tuning

Feature tuning in the change detection task was assessed using ROC analysis (Green and Swets, 1966), which quantifies how well an external observer could discriminate between two sets of values. The area under the ROC curve (AUC) was computed for the distributions of either the firing rate response of V1 neurons or video ME (0-200 ms) between grating orientations A/B or C/D, or auditory frequencies A/B or C/D. Each class had to have at least 10 trials. AUC values are in the range of 0 to 1, but were rescaled between -1 and 1, where -1 indicates complete selectivity to A/B and 1 to C/D.

Orientation and frequency tuning in the stimulus detection paradigm (Ext. Data Fig. 4) was assessed using the global Orientation Selectivity Index (gOSI). This measure adequately captures tuning in a circular domain (Ringach et al., 2002). As the stimulus set in both the visual and auditory domain was circular (visual orientations, and auditory tones due to the Shepard harmonic weights, see above) this measure captured selectivity to stimuli in both modalities similarly. The gOSI was computed as:

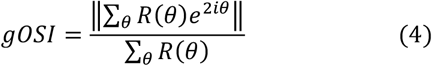

Here *R(θ)* is the firing rate response of a neuron (0-200 ms) to either a grating moving along direction *θ* or a Shepard tone with center frequency *θ* and *i* is the imaginary unit. gOSI varies between 0 and 1, with 0 indicating a neuron completely untuned, and 1 a neuron only responding to a single orientation/frequency. Neurons were deemed significantly tuned if their gOSI exceeded 95% of the shuffled distribution (recomputing gOSI for n=1000 shuffles of orientation or frequency labels). Signal correlations were computed as the Pearson correlation of the trial-averaged tuning curve between pairwise tuned neurons.

### Local field potential (LFP) analyses

The LFP was obtained by down-sampling the recorded voltage signal over time from 32000 Hz to 1024 Hz and low-pass filtered below 300 Hz (4^th^ order Butterworth filter). For current source density (CSD) and event-related potential (ERP) analyses the signal was further low-pass filtered below 100 Hz (4^th^ order Butterworth filter). The CSD profile was computed by applying standard Nicholson-Freeman calculations on the LFP signal with Vaknin transform (Vaknin et al., 1988) with 0.4 Siemens per meter as conductivity (Logothetis et al., 2007). We calculated the CSD profile for each of the linear arrays of electrodes on our polytrode configuration separately, interpolated between sites, and then merged the profiles. The ERP was the stimulus-onset locked trial-average LFP response.

To separate the sensory and motor contributions, trials were split into ‘still’ and ‘moving’ trials based on the amount of motor activity. The z-scored video ME was computed (0-500 ms post-stimulus change) and ‘still’ trials had z-scored video ME between -0.5 and 0.5 and ‘moving’ trials a z-scored video ME larger than 1. For each trial type (e.g. Amax trials changing to set A/B) the CSD and ERP was computed for still and moving trials separately. Only conditions with at least 3 trials in both still and moving conditions were included. Subsequently, the different auditory and visual trial types (saliencies and features) were averaged within modality.

We excluded a subset of sessions in which movement artefacts were present (N=16 sessions excluded from 62 sessions with video and LFP recordings in V1). To identify movement artefacts, the ERP and the CSD were computed aligned to one lick event. Sessions with movement artefacts were easily identified by the presence of low-frequency large deflections in the LFP that were several fold larger than spontaneous activity, strikingly dissimilar between adjacent channels instead of smoothly varying across cortical depth. These sessions were not included in LFP analyses.

### Cortical depth estimation

The laminar depth of each silicon probe in V1 was estimated based on a combination of two factors. First, we computed the CSD profile to contrast-reversing checkerboard stimuli. Before each session, we displayed full-field contrast-reversing checkerboards (full contrast, spatial frequency = 10 retinal degrees, temporal frequency of contrast reversal = 0.5 Hz, n=10 reversals). The earliest visible current sink was taken to indicate layer 4 (Niell and Stryker, 2008; Schnabel et al., 2018). Second, we computed the power in the 500-5000 Hz range of the raw, unfiltered signal for each channel (Senzai et al., 2019) and set the channel with highest MUA spiking power as the center of L5 at 600 μm from the dura and rereferenced all channels to this depth. Channels that were above 0 μm or below 1000 μm were excluded from the analyses.

### Regression model

To quantify single neuron encoding of sensory and motor variables we constructed a linear regression model. This approach is particularly useful to disentangle the time-dependent contribution of experimenter-controlled task events and self-timed behavioral events to single-trial neuron firing rate. The model was trained to predict the firing rate (−500 to +1000 ms relative to stimulus change in 20 ms time bins, convolved with a gaussian with a standard deviation of 25 ms). We included four sets of predictors: trial number, visual, auditory and motor variables.

The trial number predictor consisted of a whole-trial value scaled by trial number within that session. For sensory variables a separate predictor set was made per combination of orientation (or frequency) and amount of change, taking simultaneously into account the selectivity of neurons for features and saliency. For a given stimulus, there was a separate predictor for each post-stimulus time bin (50 time bins from 0 to 1000 ms). This resulted in 50 time bins x 2 saliencies (thr and max) x 2 stimuli (set A/B and set C/D) = 200 predictors per modality. For motor variables the first 25 video PCs were included. For convenience, all predictors were normalized to their maximum values before being fed into the model.

This resulted in a predictor matrix of size P x T for each neuron, where P is the number of predictors (1 trial number, 200 visual, 200 auditory, and 25 video predictors = 426 predictors) and T is the number of total time bins. The regression model was fit on concatenated single trials and T is therefore the number of trials (typically 200-500 trials per session) multiplied by the number of time bins per trial (75 time bins; -0.5 to +1 sec relative to stimulus change, 20 ms time bins).

The model was fit on catch, auditory, and visual trials (audiovisual trials were excluded during fitting) from all sessions with V1 or AC recordings during the change detection task with recorded video and without pharmacological or lick-spout manipulations. Sparsely firing neurons produced fitting difficulties and neurons with a session-average firing rate below 0.5 Hz were excluded. The total dataset for the regression analysis consisted of: n=51 sessions, NE: 9, UST: 10, MST: 32 sessions, 19217 trials, 790 V1 and 99 AC neurons. For analyses of conflict trials, the model was fit on trials excluding conflict trials and tested on the held-out conflict trials. As conflict trials were only presented in a subset of sessions and not in UST mice this dataset consisted of 37 sessions (NE: 9; MST: 28 sessions, 16021 trials, 648 V1 neurons). The model was fit with a Gaussian link function with the *glmnet* package in Matlab (Friedman et al., 2010). We used elastic-net regularization (α = 0.95) and 5-fold cross-validation. To maximally punish weights without losing model fit quality, lambda was maximized while minimizing the cross-validated error. We quantified model performance by assessing the 5-fold cross-validated explained variance (EV):

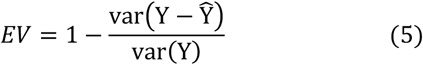

where Y is the original firing rate and Ŷ the predicted firing rate. EV was computed for the concatenated firing rate during a specified time window (0-200 ms for Figs. 3 and 4; and 0-500 ms for Fig. 5).

After fitting, the model could be used to predict firing rates on held-out test trials or with restricted predictors. To estimate the contribution of different predictors to firing rate variability, EV was computed using only one set of predictors (all other predictors were set to zero). Regularization and cross-validation already minimized overfitting to predictors, but to further verify that predictors were not capturing unrelated variance, we fit the model with one set of predictors circularly shuffled across time within the session. Thus, the original temporal relationship between for example auditory stimulus predictors and actual auditory-evoked activity was destroyed. The additional EV explained by the intact model relative to the shuffled model was taken as uniquely explained variance (Musall et al., 2019).

We found very similar results for a linear model that used smooth temporal basis functions instead of boxcar bins, or when using different elastic net mixing parameters. Note that our measure of predicting single trial binned spike counts leads to low levels of explained variance (Steinmetz et al., 2019), while we obtained high levels of explained variance when predicting trial-averaged activity (Runyan et al., 2017). Here, we were however interested in explaining single trial firing rate due to trial-by-trial differences in orofacial movements.

### Multivariate stimulus decoding from neural and video data

We used multivariate analyses to decode visual orientation or auditory frequency from either V1 population spiking activity or dominant orofacial movements (video PCs). In all decoding analyses, nearby orientations/frequencies were grouped together (denoted as set A/B versus C/D) in order to have a two-class classification problem, and here we considered only large stimulus changes (Amax and Vmax). Only sessions with V1 recordings and at least 15 trials for each orientation/frequency were included. We balanced the two classes with random subsampling of the majority class.

For the analysis in Figure 2j-o we decoded auditory or visual feature identity from simultaneously acquired data from individual sessions. We subselected all sessions which contained at least 5 neurons recorded in V1. Spikes were binned in 200 ms time bins and advanced by 25 ms. For orientation/frequency decoding using video footage, ‘population’ data was created by replacing the binned spike counts with the binned values of the first 30 principle components of the motion energy (video PCs).

We trained a support vector machine (SVM; linear kernel) using stochastic gradient descent (as implemented in scikit-learn (Pedregosa et al., 2011)) to predict the orientation/frequency at every time point. We employed 3 repeats of a 3-fold stratified cross-validation routine, whereby trials for training and testing are drawn randomly, but the equal ratio of the two classes is preserved in each set. Note that the same train/test splits are used across all time points in a given bootstrap iteration. Features (neuronal spike counts or video PCs) were standardized to have zero mean and unit variance (features of the test set were standardized using the mean and standard deviation of the training set). Reported decoding performance is the accuracy on the held-out test data. The average decoding accuracy averaged across the time bins whose edges did not exceed the 0-300 ms range were used to generate the scatter plots.

For the analyses in Figure 5g-j pseudopopulation data was constructed by combining data acquired during different sessions. We employed a bootstrapping procedure in which at every iteration we randomly sampled a subset of V1 neurons or video PCs across all recording sessions. When training the model on unisensory trials and testing on audiovisual trials (or vice versa), we selected only sessions in which for each trial type there were at least 10 trials for each orientation/frequency. When training and testing the model on the same trial type, we required at least 20 trials for each orientation/frequency. For every bootstrap iteration, we constructed a train and a test set by randomly sampling for every session 10 trials of each class for the train set, and 10 trials of each class for the test set (in the case where we trained and tested on the same trial type, trials appearing in the training set did not appear in the test set). Spikes were binned in 100 ms time bins and advanced by 25 ms. For every time point, we then assembled a feature matrix *X_t* of size 20 (2×10) by N, where N is the number of neurons, such that column *i* is the binned spike count of neuron *i* at time t in the 20 subsampled trials, and row *j* is the binned spike count at time *t* of all randomly selected neurons of a pseudo-trial that merges data from different sessions. For orientation/frequency decoding using video footage, pseudopopulation data was created in the same way by replacing the binned spike counts with the binned values of the first 30 principle components of the motion energy (video PCs). In other words, a similar data matrix was constructed subsampling video PCs (from the first 30 video PCs) during trials of specific orientation/frequency from different sessions.

The SVM was then fitted on the train set and evaluated on the test set at every time point (without a full cross-validation routine). The SVM coefficients (Fig. 5k) were obtained with an adaptation of the pseudopopulation method. For every bootstrap iteration, a random subset of 50 V1 neurons was selected. Using 3×3 stratified cross-validation routine, a linear SVM was fitted on the train set (consisting of either visual or audiovisual trials), and the coefficient of each neuron was recorded. The same pseudopopulation approach was also employed in Fig. 5g, where the orientation/frequency label was replaced by the trial type (visual versus auditory).

### Statistical analysis

Unless specified otherwise, all statistics were performed using linear mixed models (LMMs) in MatLab (MathWorks, Natick, MA). LMMs can account for the hierarchical nature of our data (neurons and trials sampled from the same mice)(Aarts et al., 2014). LMMs describe the relationship between a response variable and multiple explanatory variables, and comprise two types of explanatory terms. Fixed effects are the variables of interest, while random effects, also commonly referred to as “grouping variables”, specify and account for the group. For all analysis involving hierarchical data, LMMs were constructed with mouse identity as a random effect (intercept only). Importantly, mouse identity was not included as a random effect for analyses with cohort as fixed effect, as variability between mice was key to those results. Statistical tests were performed on the fixed effect using ANOVAs on the LMMs. To estimate the denominator degrees of freedom (DF2) for F-tests, the Satterthwaite approximation was used for LMMs. Linear hypothesis tests were performed in the case of posthoc comparisons using the relevant contrasts. Non-nested data was tested using nonparametric methods. Results with a p-value lower than 0.05 were considered significant. When multiple, independent comparisons were performed, p-values were corrected by applying a Bonferroni correction.

## EXTENDED DATA FIGURES

**Extended Data Figure 1:**
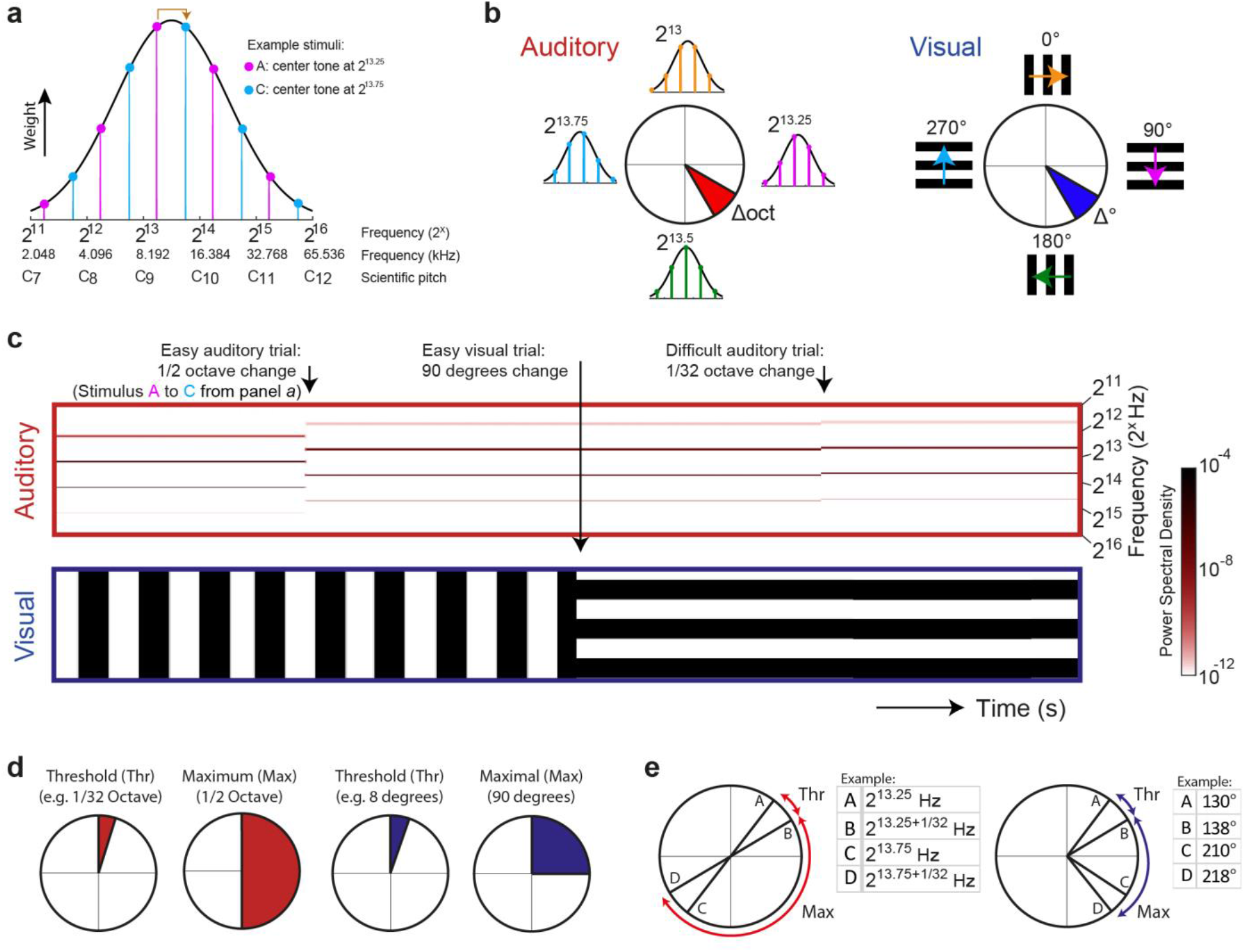
Details of auditory and visual stimulus design. **a)** Each auditory stimulus was composed of five pure tones at harmonic frequencies (octaves below and above other tones). The weight with which each tone contributed to the overall stimulus was taken from a Gaussian distribution across all possible tones. The example stimulus A in pink is composed of a tone of 2^13.25^ Hz (center tone, highest weight) and two lower (at 2^11.25^ and 2^12.25^ Hz) and two higher harmonics (at 2^14.25^ and 2^15.25^ Hz). Tones followed scientific pitch and are expressed as powers of two: 2^13^ corresponds to 8.192 kHz, and C_9_ in scientific pitch notation. During an auditory trial, the stimulus changed to a stimulus of five new harmonic tones with different weights (for example stimulus A to B). **b)** The left polar diagram shows the circular arrangement of auditory stimuli. For each cardinal direction the insets show the tonal weights associated with these stimuli. Note how ever increasing the center tone frequency ultimately results in a circular shift back to the starting stimulus. This circularity can also be seen in panel *a*: going up and down half an octave from stimulus A always results in stimulus B. The auditory stimulus set is therefore circular. This feature is exploited in the Shepard illusion of eternal rise or drop in pitch. However, our stimuli were static so the illusory effect of continuously increasing or decreasing pitch was absent. The only illusory component was that half an octave change could be both experienced as an increase or decrease in pitch. This circular design of auditory stimuli mirrors the visual stimulus set (right part) with drifting gratings in all orientations. The amount of frequency change (expressed in partial octaves, red) or orientation change (expressed in degrees, blue) determined the saliency of auditory and visual changes. **c)** Example stimuli during three consecutive trials. The upper spectrogram over time includes two auditory change trials. Auditory stimuli continued to be presented until the next auditory change, which could be identified based on a difference in spectral content, and experienced as a change in pitch. The example shows an easy auditory trial (salient change; stimulus A to B, half an octave) followed later by a difficult trial (subtle change; 1/32 of an octave). The lower schematic shows visual orientation over time including a visual trial. Note that the gratings were continuously drifting in the direction orthogonal to the grating orientation. An audiovisual trial would consist of a simultaneous change in both modalities (not shown). Note that this is only a schematic depiction, hence time is depicted in arbitrary units. **d)** Schematic of the different levels of saliency (i.e. amount of change) between threshold and maximal saliency. Threshold saliency was titrated per mouse based on task performance. **e)** The stimulus set during recording sessions was limited to four visual and four auditory stimuli with two levels of change between them. Tables show one example stimulus set for each modality, but stimuli were varied across sessions.

**Extended Data Figure 2:**
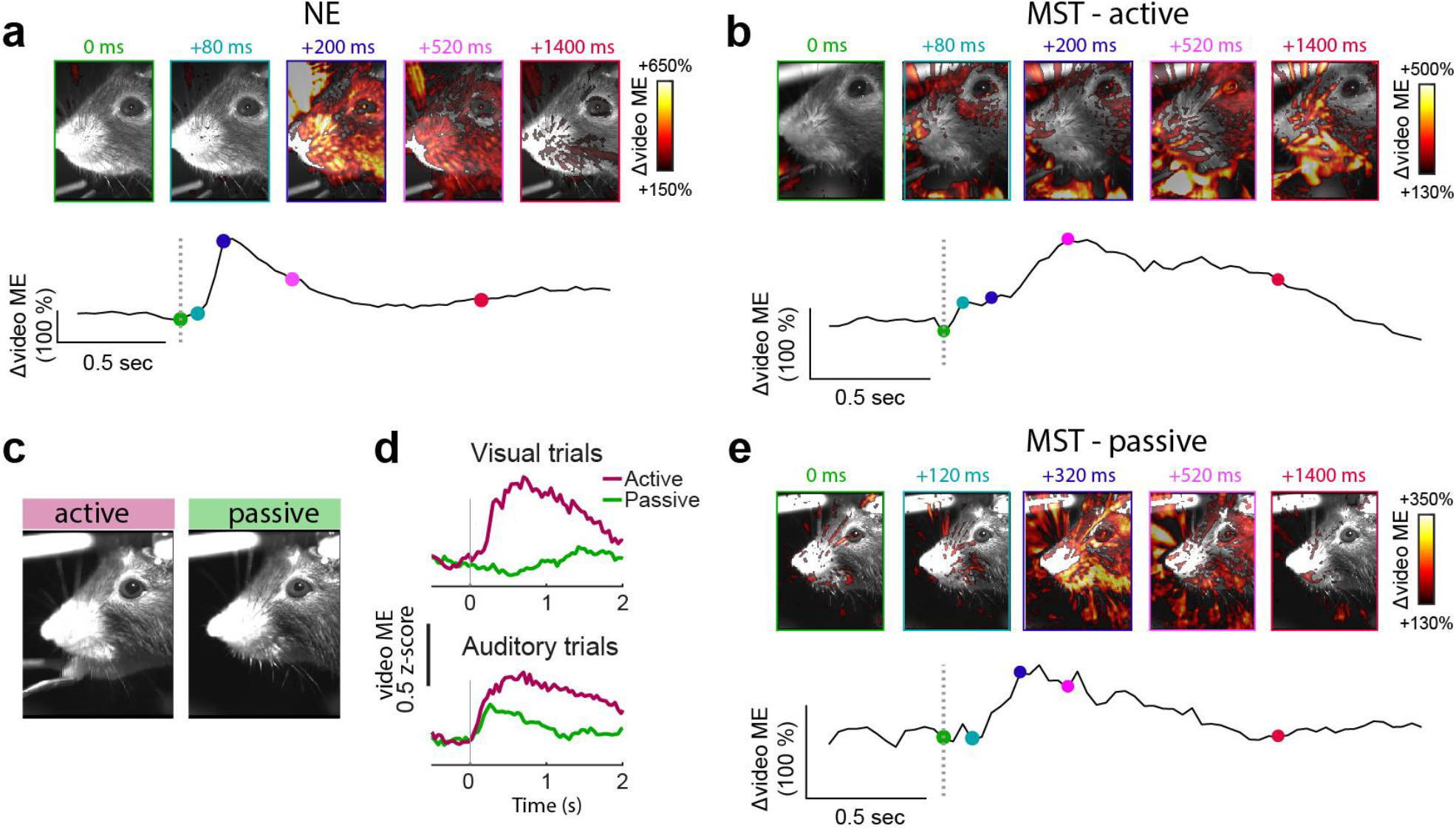
Sounds evoke instructed and uninstructed orofacial movements. **a)** Sounds evoke brief whisking and eye twitching movements in NE mice (example session). Upper images show heatmap of the increase in video ME overlaid on one reference frame. Lower trace shows video ME averaged over auditory trials with dots highlighting time points of upper frames. **b)** Same as b, but for an example MST session. Here auditory trials not only evoked whisking and eye movements (uninstructed), but also continued instrumental licking movements as mice were rewarded for reporting auditory stimuli. **c)** To further test whether the increase in motor activity was not associated with licking behavior, we continued sensory stimuli but removed the lick spout. Blocks of active trials (with lick spout, left image) and passive trials (without lick spout) were interleaved during a session. **d)** The increase in video ME normally seen following visual stimuli (due to report-related licking movements) was absent during passive blocks. On the other hand, auditory stimuli continued to evoke orofacial movements during passive blocks in the absence of licking to a rewarded lick spout. These results are in line with the comparison between cohorts (Fig. 2b) where unrewarded auditory stimuli (but not unrewarded visual stimuli) still evoke orofacial movements. Motor-related confounds are thus important to control for not only in auditory behavioral tasks, but also naive animals. **e)** Same as (a, b), but for auditory trials during passive blocks of an example MST session. Auditory trials continued to evoke uninstructed orofacial movements, but less prolonged due to the absence of licking movements.

**Extended Data Figure 3:**
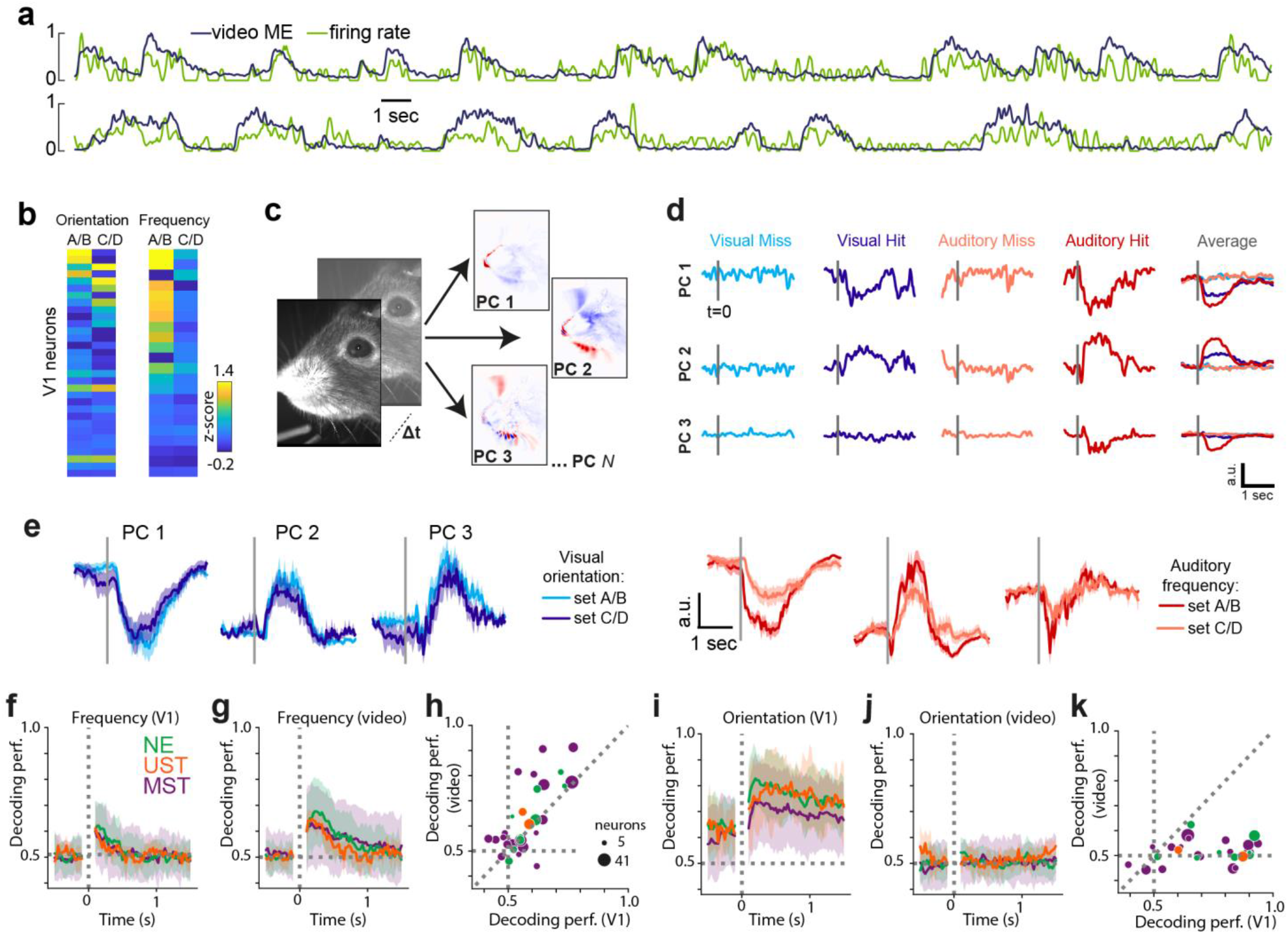
Detailed orofacial movements underlie frequency-tuned activity. **a)** Normalized firing rate and video ME over time for two example V1 neurons. Top: r=0.61, UST mouse. Bottom: r=0.55, MST mouse. **b)** Activity heatmap for simultaneously recorded V1 neurons showing a distribution of selectivity to orientations but similar tuning to auditory frequency. Left and right are taken from different sessions. **c)** To extract more detailed video information, we applied PCA to the frame-to-frame pixel intensity difference (FaceMap; Stringer et al., 2019) and extracted principal components that captured the most dominant movements (video PCs). Most movement was confined to snout, whisker pad and tongue regions. PC: principal component. **d)** Example traces of the first three PCs during individual trials of different modalities and decisions. Hit trials were associated with motor activity during lick responses and reward consumption. Data from one MST session. Gray line indicates stimulus change. **e)** First three video PCs for an example session showing similar movements following changes in visual grating orientation, but variable movements following different auditory stimuli. V1 could still encode auditory features beyond what is explained by the modulatory effects of orofacial movements. We therefore tested how well we could decode stimulus identity by considering population spiking activity in V1, and compared this to detailed video analysis. A population decoder (support vector machine, SVM) was trained to discriminate auditory or visual stimulus identity using either the spiking data or these video PC values. **f)** Auditory stimulus frequency could be decoded from V1 population activity. Decoding performance of decoders trained to discriminate post-change auditory frequency from V1 population activity. Horizontal dashed line indicates chance level. Line and shading indicate mean and 95% CI. **g)** Same as (f), but for decoding auditory frequency from the first 30 video PCs. Auditory stimulus frequency could be decoded from video data. **h)** Relationship across sessions between auditory frequency decoding performance using V1 data (x-axis) and video data (y-axis). Decoding performance was highly variable across sessions and, interestingly, strongly correlated between spiking and motor activity (R=0.71, F(1,17)=29.13, p=4.49*10^−5^). Those sessions with frequency-selective orofacial movements thus also displayed frequency-selective population activity. Further, video decoding outperformed neural decoding (F(1,49)=8.25, p=0.006). Dot size scales with number of simultaneously recorded neurons for that session and dot color indicates cohort. **i)** Same as (f), but for decoders to discriminate post-change visual orientation. Visual grating orientation could be decoded from V1 population activity. Baseline coding results from the fact that gratings jumped between the same stimuli (A/B to C/D and vice versa) and neurons showed persistent selectivity, seen in (Fig. 2h). **j)** Same as (i), but for visual orientation. Visual grating orientation could not be decoded from video PCs. **k)** Same as (h), but for the relationship between orientation decoding based on V1 activity versus video PCs. Decoding performance was not correlated across sessions (R=0.15,F(1,28)=0.67, p=0.42) and higher for V1 spikes than for video PCs (F(1,42)=51.14, p=9.09*10^−9^). Although absolute decoding performance from these qualitatively different sources is less meaningful, the dissimilarity between modalities is striking.

**Extended Data Figure 4:**
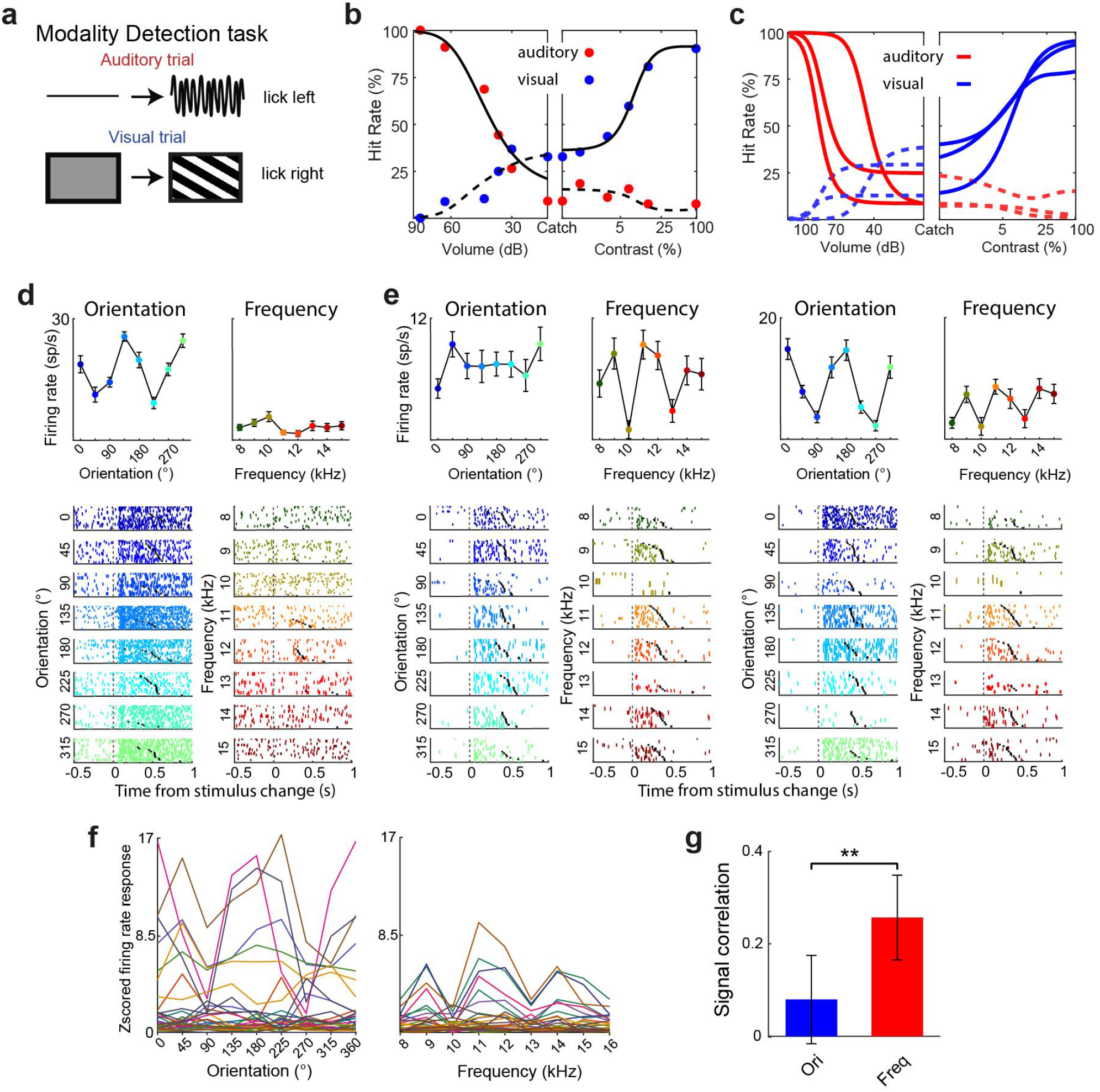
Similar frequency tuning of primary visual cortical neurons during audiovisual stimulus detection. **a)** To establish whether our findings generalized beyond our change detection task, we trained animals (n=3) to detect the presence of auditory and visual stimuli (same stimulus set as in the change detection task) and to discriminate and selectively report the modality, as in the MST task of our main change detection paradigm. Rewards were allocated upon licking to the auditory lick spout after the onset of one of eight tones, and upon licking to the visual lick spout to one of eight gratings was rewarded. **b)** Performance on an example session on the detection of auditory stimuli of varying volume (left panel) and of varying contrast (right panel). Note how auditory and visual hit rates increase as a function of volume and contrast, respectively. The behavioral data was fit with the same two-alternative signal detection model as behavioral data from the change detection task. Behavioral response rates are shown as dots, model fits as lines. **c)** Average psychometric fits for each mouse obtained by averaging the parameters of single session fits. **d)** Raster plot and tuning curve of an example orientation-tuned V1 neuron. Upper panels show firing rate (0-200 ms) in response to eight drifting grating orientations (left) and eight compound Shepard tones with center tone spaced between 8 and 15 kHz (right). Dot and error bar show mean ± SEM across trials. Colored tickmarks in the lower raster plots show trial-by-trial spiking. Black tick marks indicate first lick after the stimulus. Note the classical orientation tuning expected from V1 neurons in response to full-field oriented drifting gratings. Auditory frequency tuning was not significant. **e)** Same as (d), but for two V1 neurons from the same session where the auditory response depended on the frequency components of the auditory stimulus. Note how the neurons are similarly tuned and their firing rates are associated with licking behavior as well. **f)** Tuning curves for orientation and frequency for all V1 neurons (individual lines) from one session. Note dissimilarity in orientation tuning, but similarity in frequency tuning. **g)** The signal correlation of all significantly orientation-tuned (left) and frequency-tuned (right) neurons. Signal correlations were computed as the Pearson correlation of trial-averaged tuning curves between neuronal pairs. Signal correlation was higher between frequency-tuned neurons than orientation-tuned neurons (F(1,406)=9.50, p=0.0022; n=148 signal correlations from 23 orientation-tuned V1 neurons, n=258 from 36 frequency-tuned V1 neurons). The finding that V1 neurons responded to the same frequencies (those associated with motor movement, Fig. 2i; Ext. Data Fig. 3b) suggests that variability in motor variables drives tuning. **p<0.01.

**Extended Data Figure 5:**
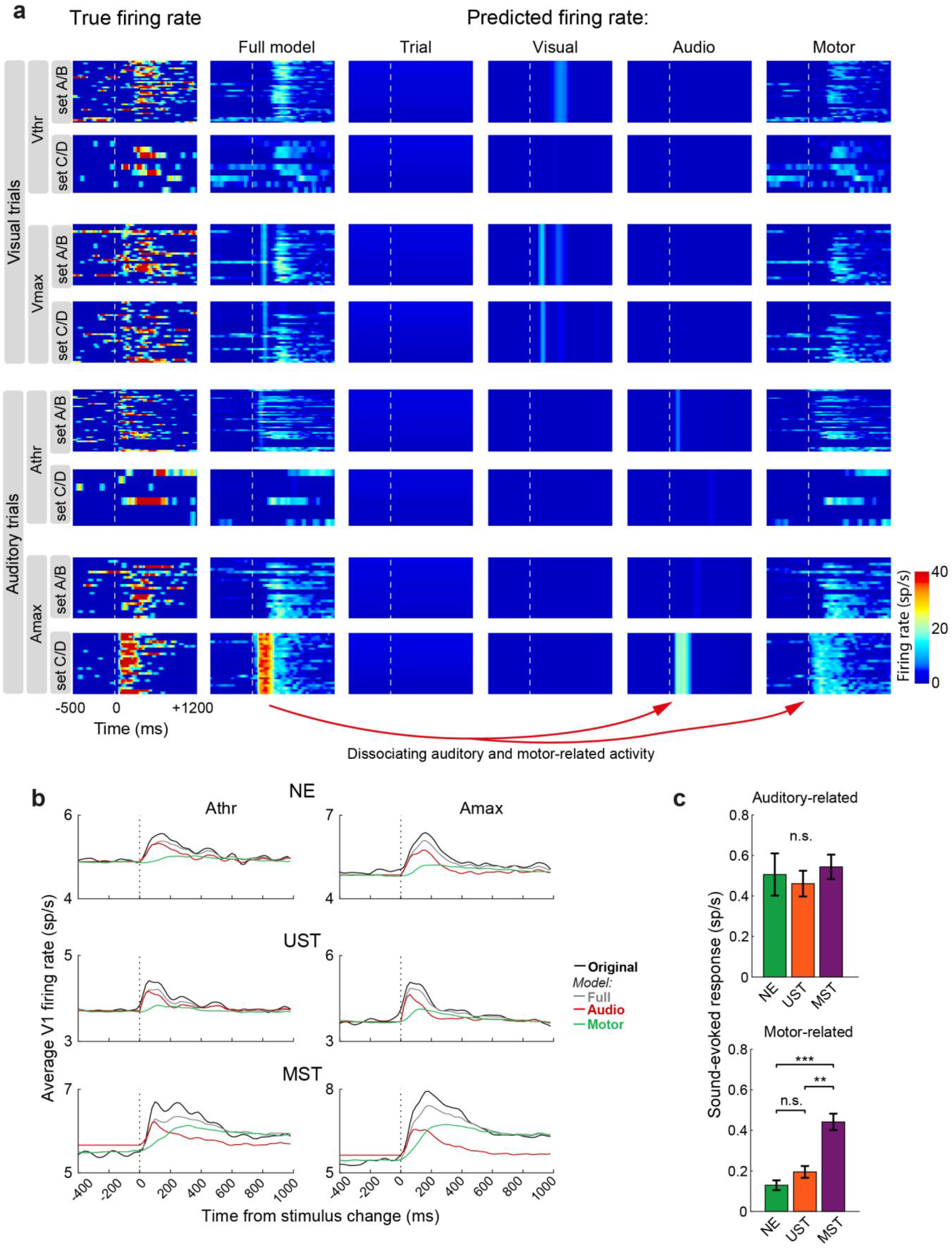
Dissociating visual, auditory and motor-related activity using a regression model. **a)** Each heatmap shows the firing rate over time for a subset of trials with each row representing a different trial type, and each column a different source of the firing rate. The leftmost column shows the original firing rate. The second column shows the predicted firing rate for the same trials using all predictors in the model. The remaining columns show the predicted firing rate using only a subset of the predictors. For this example neuron, the trial number explained little variability (trial number captured response drift across the session for some other neurons, not shown). Visual predictors explained an early response transient especially in Vmax trials. Auditory predictors captured an early response transient in some auditory trials (set C/D), whereas motor variables (the first 25 video PCs) captured variability across visual and auditory trial types. **b)** Same as Figure 3f, but for each of the task cohorts separately and auditory trials only. Auditory-related activity was present in all three cohorts. Sound-evoked motor-related activity was larger in the MST cohort, quantified in (c). **c)** Predicted sound-evoked response (0-200 ms minus baseline activity) for each of the cohorts using either auditory predictors (top) or motor predictors only (bottom). Cohorts did not significantly differ in auditory-related activity (F(2,790)=0.18, p=0.835), while motor-related activity was significantly different (F(2,790)=8.07, p=0.00034) and significantly larger in MST mice compared to NE and UST (Posthoc comparison: NE vs. MST: F(1,787)=11.4, p=0.000789; UST vs. MST: F(1,787)=7.3, p=0.00702; NE vs. UST: F(1,787)=0.3, p=0.582). The larger sound-evoked response in MST mice in Figure 1e,f is therefore attributable to increased motor-related and not auditory-related activity. Mean ± SEM across neurons.

**Extended Data Figure 6:**
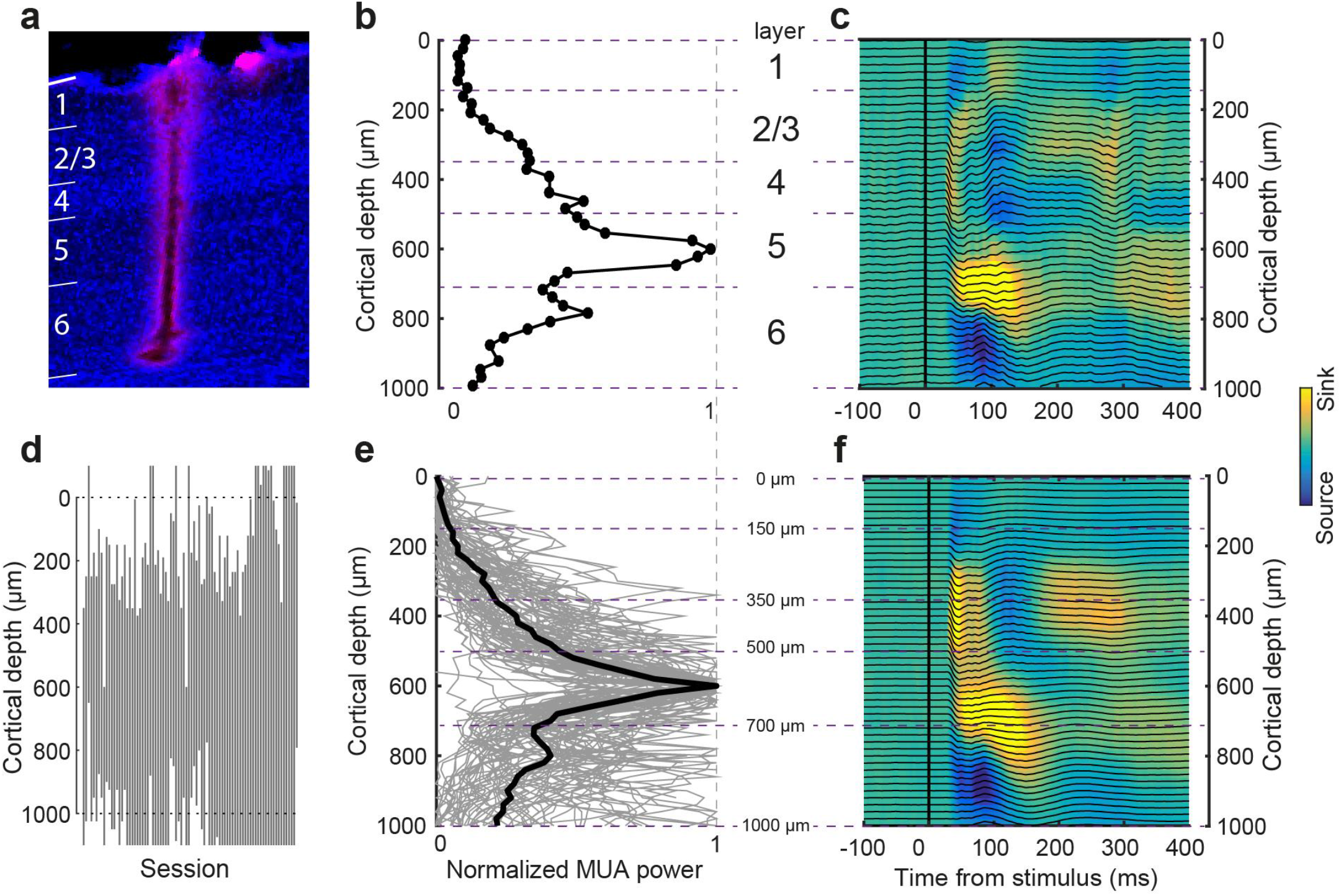
Cortical depth estimation in primary visual cortex using physiological markers. **a)** Close-up of a coronal section on V1 showing the electrode track stained with DiI. **b)** Example distribution along the probe of spectral power (500 Hz to 5 kHz) indicative of multi-unit activity (MUA). High MUA power is characteristic of L5. Compare with (Senzai et al., 2019). **c)** Current source density (CSD) map and LFP traces (black lines) in response to checkerboard stimulation. *B* and *C* are from the same example session. Color corresponds to CSD power. **d)** Overview of electrode span across layers. Each line is one session (n=84 sessions). Data from electrodes at depths above 0 or below 1000 μm were excluded from analyses. **e)** Same as b, but for all sessions. Gray lines are individual sessions, black line the median. **f)** Same as c, but averaged across all sessions.

**Extended Data Figure 7:**
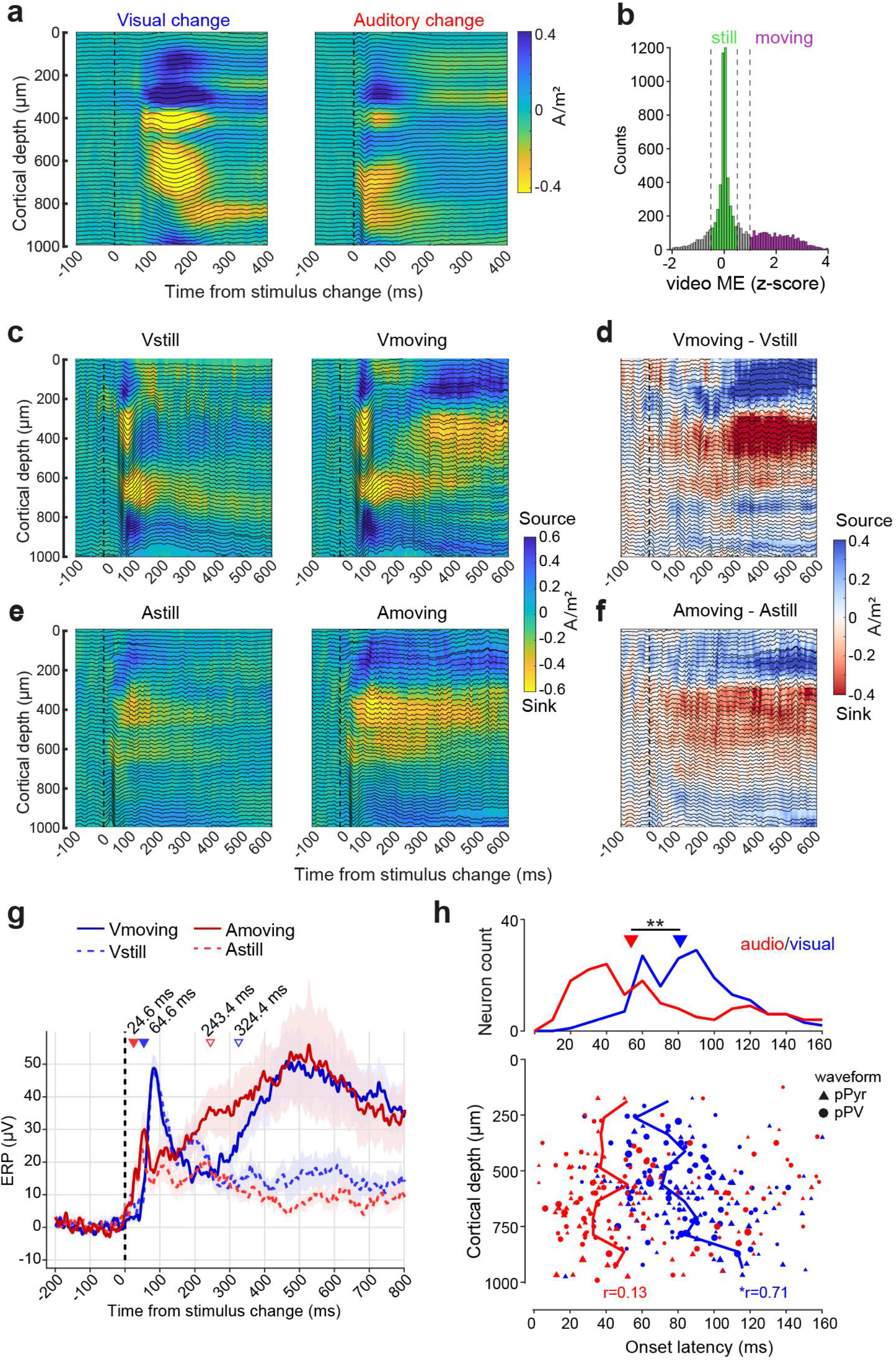
Early sensory and late motor-related components of current-source density and cell-type spiking profiles in visual cortex. **a)** The current source density (colormap, CSD) and event-related potential (black traces, ERP) for auditory and visual stimulus changes in the same example session (MST mouse). **b)** Histogram of z-scored video ME (0-500 ms post-change) across visual and auditory trials of all sessions with LFP recordings in V1 (all cohorts). To separate the contribution of motor activity to the LFP, all trials were split into ‘still’ and ‘moving’ trials based on the amount of motor activity. ‘Still’ trials had z-scored video ME between -0.5 and 0.5 and ‘moving’ trials a z-scored video ME larger than 1. **c)** For each session a CSD map was constructed using either still or moving trials given the same visual stimuli. Average across n=46 sessions (NE: 12 sessions; UST: 7; MST: 27). Visual stimuli evoked a consistent and characteristic current source density (CSD) profile with an early sink in L4 and subsequent sink-source pairs in L2/3 and L5/6, in line with earlier reports (Niell and Stryker, 2008; Schnabel et al., 2018; Senzai et al., 2019). **d)** The difference between the Vstill and Vmoving maps in (c), which we interpret as mostly related to motor differences. Note how most of the motor-related CSD power is expressed after 200 ms in L2-5 and predominantly in superficial and middle layers. **e)** Same as (c), but for auditory trials. Note how the early sinks and sources in deep layers of the auditory CSD map in the example session of (a) are only partially reflected in the average. **(f)** Difference map of the Astill and Amoving maps in (e). Note how the movement-associated CSD pattern resembles that of visual trials (d), but is generated somewhat earlier in time. **g)** Absolute ERP response (in μV) averaged across cortical depth for selected trial categories. The tick marks and text denote the first time bin the LFP response is different from baseline (−500 to 0 ms) during auditory or visual trials irrespective of motor activity (p<0.05, Wilcoxon signed rank test, Bonferroni correction). These latencies closely match spiking onset latencies (Fig. 3b). The LFP response for auditory trials can be seen to diverge between still and moving trials around 100 ms after stimulus onset and was significantly different after 243.4 ms (bootstrap test, n=1000 resamples, p<0.05) and after 324.4 ms for visual trials (p<0.05) suggestive of late motor-related signals. Line and shading are mean ± SEM. **h)** Laminar organization of onset latencies of visual and auditory responses in V1 (spiking data, not LFP). Top histogram shows the distribution of onset latencies of all significantly auditory responsive neurons (red) and visually responsive neurons (blue). Significance and onset latency were assessed using a binning-free algorithm, ZETA (Montijn et al., 2021). Spiking onset was significantly earlier for auditory versus visual stimuli (55.3 ms (31.4 - 108.5 ms) versus 80.3 ms (61.5 - 98.5 ms); median and interquartile range; F(1,411)=5.37, p=0.0209), similar to our earlier population-averaged approach (Fig. 3b). Bottom panel shows each neuron’s onset latency as a function of its recorded depth and cell type. If neurons are bimodally responsive they appear twice. Symbols are scaled by response magnitude. Putative pyramidal cells (broad-spiking) and putative parvalbumin expressing cells (narrow-spiking) were classified based on their waveform. L1 is mostly empty because almost no cells were recorded in that layer. *p<0.05, **p<0.01. Visually driven cells first began to fire significantly in the middle and superficial layers and later in deeper layers, consistent with the canonical sensory processing scheme (Douglas and Martin, 2004; Harris and Shepherd, 2015). Auditory-evoked firing started at similar latencies across layers, with many auditory responsive neurons in deep layers. Cortical depth was significantly correlated to spiking onset latency during visual trials (r=0.71, p=0.015, Pearson correlation), but not auditory trials (r=0.13, p=0.696).

**Extended Data Figure 8:**
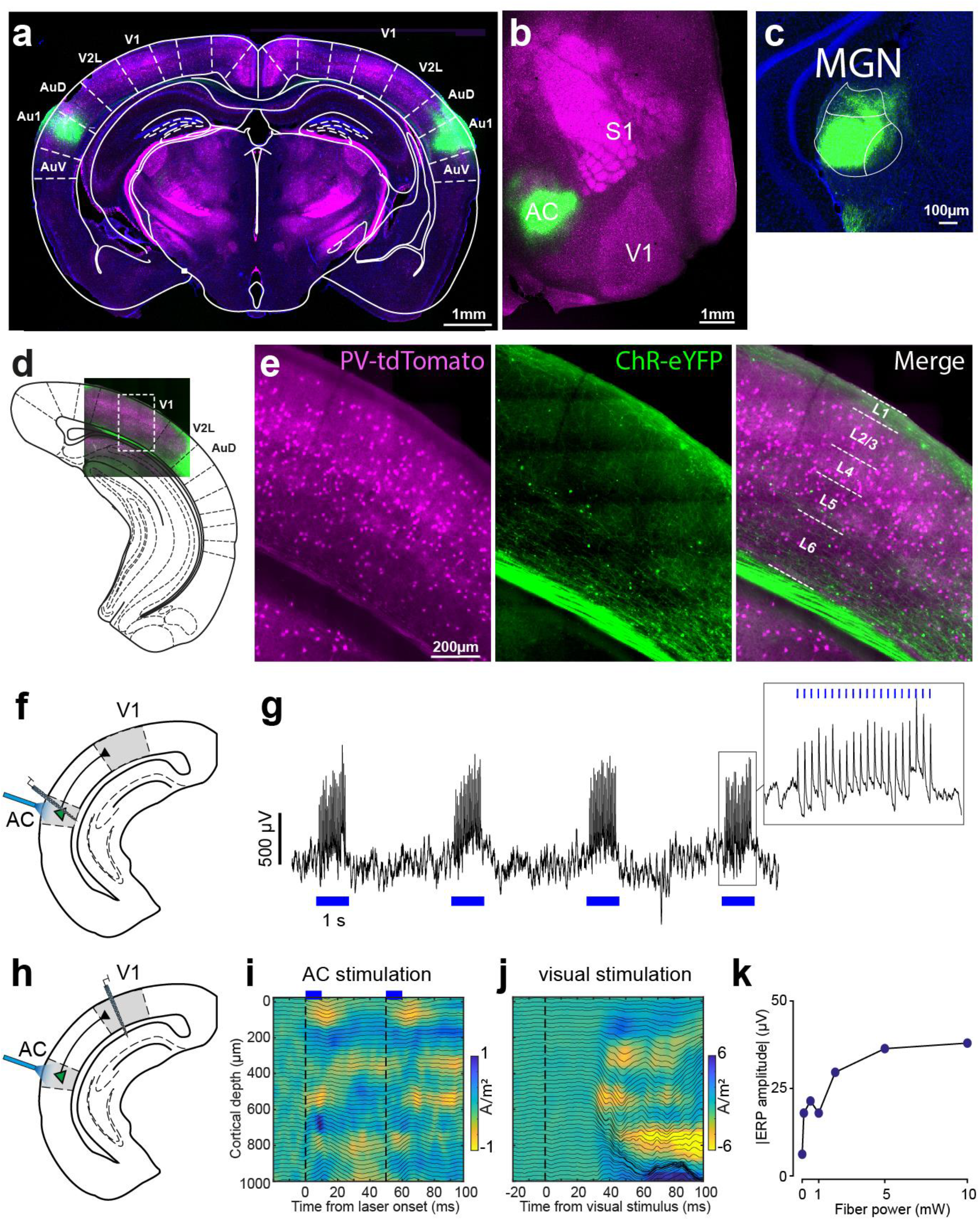
Auditory cortical projections modulate superficial and deep layers of primary visual cortex. **a)** Coronal section showing bilateral AC expression of AAV2-CaMKIIa-hChR2(H134R)-eYFP (*green*: eYFP) in a PvCre-tdTomato mouse (magenta: tdTomato), centered at primary auditory cortex (Au1). AuV: ventral secondary auditory cortex. AuD: dorsal secondary auditory cortex. V2L: lateral secondary visual cortex. **b)** Same as a, but for a flattened cortical section showing ChR2-eYFP expression in AC. **c)** Close up of densely labeled projections in medial geniculate nucleus of the thalamus (MGN) confirming infection of AC. **d)** Reference section with the box outlining the location of close up image shown in (E). **e)** Close up image of highlighted section in D showing axonal terminals in superficial L1and L5/6. **f)** Schematic of the experiment verifying optogenetic excitation of AC cell bodies. **g)** Raw voltage trace from an example electrode in AC during AC photostimulation, verifying effective optogenetic recruitment of local neurons. 5 mW, 10 ms pulses @ 20 Hz. **h)** Schematic of the experiment to optogenetically stimulate AC cell bodies and record laminar LFP in V1. **i)** CSD and LFP profile in V1 during AC photostimulation (average of n=2 sessions in 2 animals). Note how pulsed AC stimulation gives rises to a repetitive CSD response (sink) in the superficial (< 150 μm) and middle/deeper layers (500-800 μm). Vertical dashed lines indicate repeated AC stimulation. **j)** Same as in (i), but for an example visual checkerboard stimulation for comparative purposes. **k)** The event-related potential (ERP) following photopulses (+5 to +20 ms after pulse) increases as a function of fiber power, suggesting optogenetic stimulation affects V1 LFP in a dose-dependent manner. The ERP response was obtained by averaging the absolute signal from channels over all cortical depths.

**Extended Data Figure 9.**
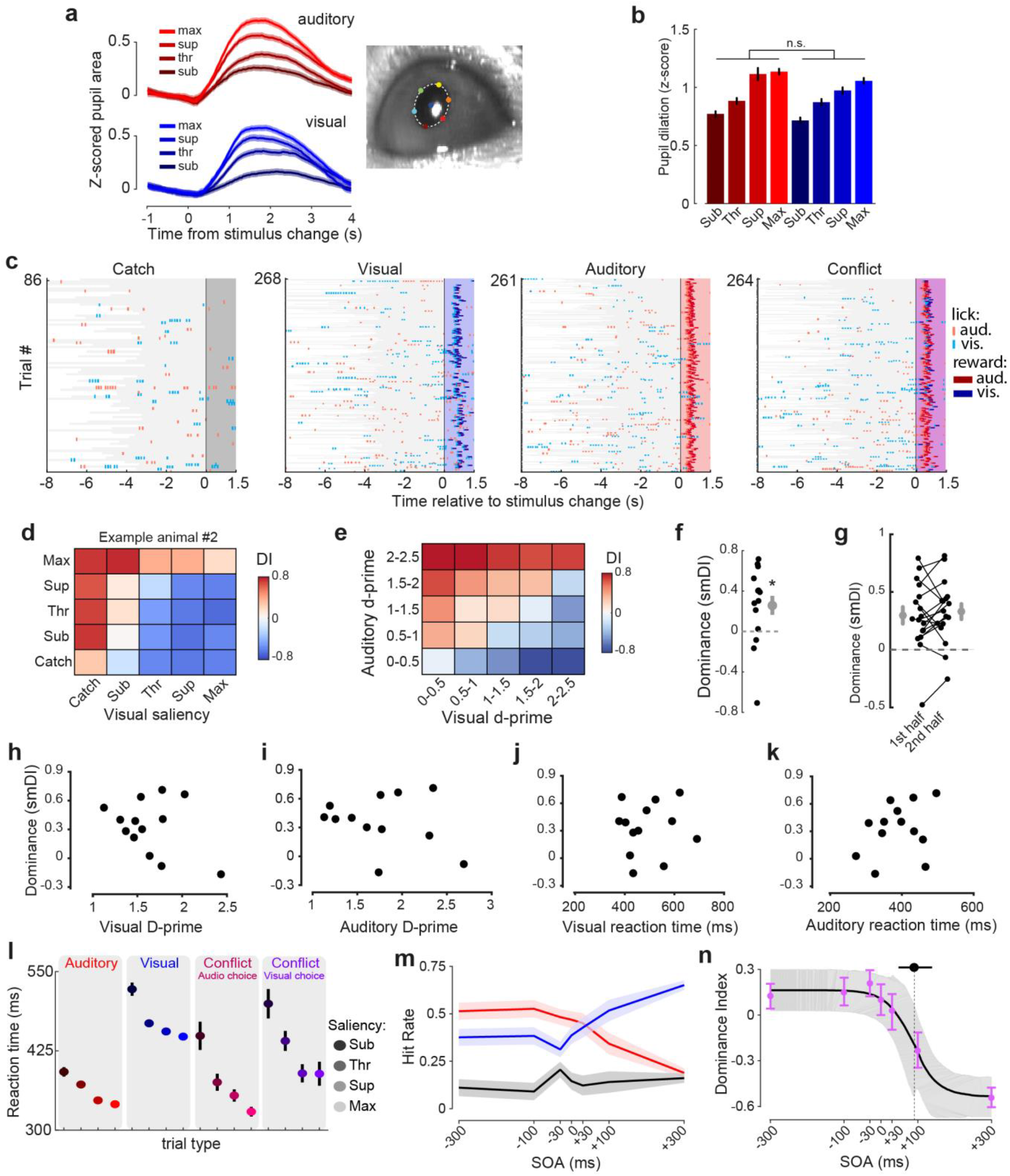
Auditory behavioral dominance in conflict trials is stable, independent of performance, and depends on relative stimulus timing. **a)** As a proxy for subjective saliency or arousal, we measured pupil dilation over time for saliency-matched auditory and visual trials. Cropped image shows pupil fit. Line and shading indicate mean ± SEM across N = 40 sessions from 9 mice. All data in this figure are from MST mice. **b)** Quantification of maximal pupil dilation. The effect of modality on pupil dilation was tested in a linear mixed model with fixed effects of hit/miss, saliency, modality and random effect of mouse ID. Whether it was a hit or miss had the largest effect (F(1,7530)=1138.95, p=1.24*10^−232^), then saliency (F(1,7524)=33.36, p=7.97*10^−9^, with no effect of modality (F(1,7526)=1.53, p=0.2164). This supports the idea that visual and auditory conditions were matched in subjective saliency. **c)** Auditory dominance in an example session. Raster plots show for each trial type licks and rewards at the auditory and visual lick spout (red and blue tick marks respectively) aligned to stimulus change (t=0). Colored zones indicate response window (0 to 1.5s). Gray: inter-trial interval. Licks before t=0 were spontaneous. Note how during conflict trials, auditory licks and rewards dominate. **d)** Dominance index (DI) heatmap (as in Fig. 5b) for the only animal out of 17 MST mice) displaying visual dominance. **e)** A heatmap of the auditory dominance index for conditions binned based on performance (d-prime) on unimodal trials. This is in contrast to the analyses presented in the main text, where conditions were grouped based on the predetermined saliency gauged by psychophysical performance in previous sessions. The current analysis controls for changes in performance by reassigning each bin of the heatmap to d-prime levels within that session. It can be seen that performance-matched conflict trial conditions (along the diagonal) have positive dominance index values, confirming auditory dominance. **f)** The saliency-matched dominance index (smDI) for conflict trials that are matched in performance to unimodal trials (conditions along the bottom-left to top-right diagonal of *e*) is significantly different from zero (Wilcoxon signed rank test, n=17 mice, p=0.030). Grey dot is mean ± SEM, *p <0.05. In (f) to (l), each dot is the smDI of one animal. **g)** Auditory dominance was stable across the session, with auditory dominance computed on the first and second half of sessions being similar (Wilcoxon signed rank test, n=17 mice, p=0.492). **h)** Dominance was not correlated with visual performance (d-prime on unimodal trials of maximal visual saliency in the same sessions; r=-0.29, p=0.26). **i)** Dominance was not correlated with auditory performance (d-prime on unimodal trials of maximal auditory saliency in the same sessions; r=0.24, p=0.35). **j)** Dominance was not correlated with mean reaction time in visual trials (r=0.38, p=0.14). **k)** Dominance was not correlated with mean reaction time in auditory trials (r=0.09, p= 0.73). **l)** Reaction times on visual, auditory and conflict trials. For conflict trials, only saliency-matched conflicts are shown (Vsub + Asub, Vthr + Athr, etc.). Conflict trials were split based on choice. Mean ± SEM. **m)** We varied stimulus onset asynchrony between auditory and visual stimulus changes during conflict trials. The plot shows the percentage of auditory choice (red), visual choice (blue) or no lick (black) during saliency-matched threshold-level conflict trials as a function of stimulus onset asynchrony (SOA). A positive SOA value means that the visual change was presented first, followed by the auditory change. **n)** Purple error bars show mean and standard deviation of DI as a function of stimulus-onset asynchrony. Black line and gray shading show bootstrapped cumulative Gaussian fit of DI as a function of SOA (median and 95% confidence interval). Top error bar and dotted line indicate crossover point, i.e. fitted *μ* parameter (median and 95% confidence interval). Auditory dominance reverses once the visual stimulus change precedes the auditory change by 89.9 ms (95% CI: 47.7-138.7 ms). This is close to the difference in reaction time between saliency matched auditory and visual conditions: 110.5 ms on average. In other words, when the visual change preceded the auditory change by about 90 ms, auditory dominance was halfway to reversing into visual dominance. Further advancing the visual change in time completely reversed the dominance. This may reflect a scenario in which the visual evidence has instructed the decision-making system to an extent that subjects have already committed to a motor plan (namely to lick the visual spout) before the auditory evidence may take control. Similar temporal dominance of audition over vision has been reported in humans (Burr et al., 2009; Repp and Penel, 2002; Shams et al., 2000).

## References

Aarts, E., Verhage, M., Veenvliet, J.V., Dolan, C.V., van der Sluis, S., 2014. A solution to dependency: using multilevel analysis to accommodate nested data. Nat. Neurosci. 17, 491–496. https://doi.org/10.1038/nn.3648

Bimbard, C., Sit, T.P., Lebedeva, A., Harris, K.D., Carandini, M., 2021. Behavioral origin of sound-evoked activity in visual cortex. BioRxiv 2021.07.01.450721. https://doi.org/10.1101/2021.07.01.450721

Bizley, J.K., Nodal, F.R., Bajo, V.M., Nelken, I., King, A.J., 2007. Physiological and Anatomical Evidence for Multisensory Interactions in Auditory Cortex. Cereb. Cortex 17, 2172– 2189. https://doi.org/10.1093/cercor/bhl128

Bos, J.J., Vinck, M., van Mourik-Donga, L.A., Jackson, J.C., Witter, M.P., Pennartz, C.M.A., 2017. Perirhinal firing patterns are sustained across large spatial segments of the task environment. Nat. Commun. 8, 15602. https://doi.org/10.1038/ncomms15602

Bouvier, G., Senzai, Y., Scanziani, M., 2020. Head Movements Control the Activity of Primary Visual Cortex in a Luminance-Dependent Manner. Neuron 108, 500-511.e5. https://doi.org/10.1016/j.neuron.2020.07.004

Budinger, E., Heil, P., Hess, A., Scheich, H., 2006. Multisensory processing via early cortical stages: Connections of the primary auditory cortical field with other sensory systems. Neuroscience 143, 1065–1083. https://doi.org/10.1016/j.neuroscience.2006.08.035

Budinger, E., Scheich, H., 2009. Anatomical connections suitable for the direct processing of neuronal information of different modalities via the rodent primary auditory cortex. Hear. Res., Multisensory integration in auditory and auditory-related areas of cortex 258, 16–27. https://doi.org/10.1016/j.heares.2009.04.021

Burr, D., Banks, M.S., Morrone, M.C., 2009. Auditory dominance over vision in the perception of interval duration. Exp. Brain Res. 198, 49. https://doi.org/10.1007/s00221-009-1933-z

Campi, K.L., Bales, K.L., Grunewald, R., Krubitzer, L., 2010. Connections of Auditory and Visual Cortex in the Prairie Vole (Microtus ochrogaster): Evidence for Multisensory Processing in Primary Sensory Areas. Cereb. Cortex 20, 89– 108. https://doi.org/10.1093/cercor/bhp082

Cappe, C., Barone, P., 2005. Heteromodal connections supporting multisensory integration at low levels of cortical processing in the monkey. Eur. J. Neurosci. 22, 2886–2902. https://doi.org/10.1111/j.1460-9568.2005.04462.x

Ceballo, S., Piwkowska, Z., Bourg, J., Daret, A., Bathellier, B., 2019. Targeted Cortical Manipulation of Auditory Perception. Neuron 104, 1168-1179.e5. https://doi.org/10.1016/j.neuron.2019.09.043

Chou, X., Fang, Q., Yan, L., Zhong, W., Peng, B., Li, H., Wei, J., Tao, H.W., Zhang, L.I., 2020. Contextual and cross-modality modulation of auditory cortical processing through pulvinar mediated suppression. eLife 9, e54157. https://doi.org/10.7554/eLife.54157

Coen, P., Sit, T.P.H., Wells, M.J., Carandini, M., Harris, K.D., 2021. Mouse frontal cortex mediates additive multisensory decisions. bioRxiv 2021.04.26.441250. https://doi.org/10.1101/2021.04.26.441250

DeWeese, M.R., Wehr, M., Zador, A.M., 2003. Binary Spiking in Auditory Cortex. J. Neurosci. 23, 7940–7949. https://doi.org/10.1523/JNEUROSCI.23-21-07940.2003

Douglas, R.J., Martin, K.A.C., 2004. Neuronal Circuits of the Neocortex. Annu. Rev. Neurosci. 27, 419–451. https://doi.org/10.1146/annurev.neuro.27.070203.144152

Falchier, A., Clavagnier, S., Barone, P., Kennedy, H., 2002. Anatomical Evidence of Multimodal Integration in Primate Striate Cortex. J. Neurosci. 22, 5749–5759.

Fetsch, C.R., DeAngelis, G.C., Angelaki, D.E., 2013. Bridging the gap between theories of sensory cue integration and the physiology of multisensory neurons. Nat. Rev. Neurosci. 14, 429–442. https://doi.org/10.1038/nrn3503

Fishman, M.C., Michael, C.R., 1973. Integration of auditory information in the cat’s visual cortex. Vision Res. 13, 1415– 1419. https://doi.org/10.1016/0042-6989(73)90002-3

Friedman, J., Hastie, T., Tibshirani, R., 2010. Regularization Paths for Generalized Linear Models via Coordinate Descent. J. Stat. Softw. 33, 1–22.

Fu, Y., Tucciarone, J.M., Espinosa, J.S., Sheng, N., Darcy, D.P., Nicoll, R.A., Huang, Z.J., Stryker, M.P., 2014. A Cortical Circuit for Gain Control by Behavioral State. Cell 156, 1139– 1152. https://doi.org/10.1016/j.cell.2014.01.050

Garner, A.R., Keller, G.B., 2022. A cortical circuit for audio-visual predictions. Nat. Neurosci. 25, 98–105. https://doi.org/10.1038/s41593-021-00974-7

Ghazanfar, A.A., Schroeder, C.E., 2006. Is neocortex essentially multisensory? Trends Cogn. Sci. 10, 278–285. https://doi.org/10.1016/j.tics.2006.04.008

Green, D.M., Swets, J.A., 1966. Signal detection theory and psychophysics, Signal detection theory and psychophysics. John Wiley, Oxford, England.

Guitchounts, G., Masís, J., Wolff, S.B.E., Cox, D., 2020. Encoding of 3D Head Orienting Movements in the Primary Visual Cortex. Neuron 108, 512-525.e4. https://doi.org/10.1016/j.neuron.2020.07.014

Harris, K.D., Shepherd, G.M.G., 2015. The neocortical circuit: themes and variations. Nat. Neurosci. 18, 170–181. https://doi.org/10.1038/nn.3917

Henry, K.R., Lepkowski, C.M., 1978. Evoked Potential Correlates of Genetic Progressive Hearing Loss:Age-related Changes from the Ear to the Inferior Colliculus ofC57BL/6 and CBA/J Mice. Acta Otolaryngol. (Stockh.) 86, 366–374. https://doi.org/10.3109/00016487809124758

Henschke, J.U., Noesselt, T., Scheich, H., Budinger, E., 2015. Possible anatomical pathways for short-latency multisensory integration processes in primary sensory cortices. Brain Struct. Funct. 220, 955–977. https://doi.org/10.1007/s00429-013-0694-4

Ibrahim Mesik, L., Ji, X., Fang, Q., Li, H., Li, Y., Zingg, B., Zhang, L.I., Tao, H.W., 2016. Cross-Modality Sharpening of Visual Cortical Processing through Layer-1-Mediated Inhibition and Disinhibition. Neuron 89, 1031–1045. https://doi.org/10.1016/j.neuron.2016.01.027

Iurilli, G., Ghezzi, D., Olcese, U., Lassi, G., Nazzaro, C., Tonini, R., Tucci, V., Benfenati, F., Medini, P., 2012. Sound-Driven Synaptic Inhibition in Primary Visual Cortex. Neuron 73, 814– 828. https://doi.org/10.1016/j.neuron.2011.12.026

Jones, E.G., Powell, T.P.S., 1970. An Anatomical Study of Converging Sensory Pathways Within the Cerebral Cortex of the Monkey. Brain 93, 793–820. https://doi.org/10.1093/brain/93.4.793

Kayser, C., Logothetis, N.K., 2007. Do early sensory cortices integrate cross-modal information? Brain Struct. Funct. 212, 121–132. https://doi.org/10.1007/s00429-007-0154-0

Knöpfel, T., Sweeney, Y., Radulescu, C.I., Zabouri, N., Doostdar, N., Clopath, C., Barnes, S.J., 2019. Audio-visual experience strengthens multisensory assemblies in adult mouse visual cortex. Nat. Commun. 10, 5684. https://doi.org/10.1038/s41467-019-13607-2

Leinweber, M., Ward, D.R., Sobczak, J.M., Attinger, A., Keller, G.B., 2017. A Sensorimotor Circuit in Mouse Cortex for Visual Flow Predictions. Neuron 95, 1420-1432.e5. https://doi.org/10.1016/j.neuron.2017.08.036

Lippert, M., Logothetis, N.K., Kayser, C., 2007. Improvement of visual contrast detection by a simultaneous sound. Brain Res. 1173, 102–109. https://doi.org/10.1016/j.brainres.2007.07.050

Logothetis, N.K., Kayser, C., Oeltermann, A., 2007. In Vivo Measurement of Cortical Impedance Spectrum in Monkeys: Implications for Signal Propagation. Neuron 55, 809–823. https://doi.org/10.1016/j.neuron.2007.07.027

Mathis, A., Mamidanna, P., Cury, K.M., Abe, T., Murthy, V.N., Mathis, M.W., Bethge, M., 2018. DeepLabCut: markerless pose estimation of user-defined body parts with deep learning. Nat. Neurosci. 21, 1281–1289. https://doi.org/10.1038/s41593-018-0209-y

Meijer, G.T., Marchesi, P., Mejias, J.F., Montijn, J.S., Lansink, C.S., Pennartz, C.M.A., 2020. Neural Correlates of Multisensory Detection Behavior: Comparison of Primary and Higher-Order Visual Cortex. Cell Rep. 31, 107636. https://doi.org/10.1016/j.celrep.2020.107636

Meijer, G.T., Mertens, P.E.C., Pennartz, C.M.A., Olcese, U., Lansink, C.S., 2019. The circuit architecture of cortical multisensory processing: Distinct functions jointly operating within a common anatomical network. Prog. Neurobiol. 174, 1–15. https://doi.org/10.1016/j.pneurobio.2019.01.004

Meijer, G.T., Montijn, J.S., Pennartz, C.M.A., Lansink, C.S., 2017. Audio-visual modulation in mouse V1 depends on cross-modal stimulus configuration and congruency. J. Neurosci. Off. J. Soc. Neurosci. https://doi.org/10.1523/JNEUROSCI.0468-17.2017

Meijer, G.T., Pie, J.L., Dolman, T.L., Pennartz, C.M.A., Lansink, C.S., 2018. Audiovisual Integration Enhances Stimulus Detection Performance in Mice. Front. Behav. Neurosci. 12. https://doi.org/10.3389/fnbeh.2018.00231

Mesik, L., Huang, J.J., Zhang, L.I., Tao, H.W., 2019. Sensory-and Motor-Related Responses of Layer 1 Neurons in the Mouse Visual Cortex. J. Neurosci. 39, 10060–10070. https://doi.org/10.1523/JNEUROSCI.1722-19.2019

Miller, M.W., Vogt, B.A., 1984. Direct connections of rat visual cortex with sensory, motor, and association cortices. J. Comp. Neurol. 226, 184–202.

Minamimoto, T., Kimura, M., 2002. Participation of the Thalamic CM-Pf Complex in Attentional Orienting. J. Neurophysiol. 87, 3090–3101. https://doi.org/10.1152/jn.2002.87.6.3090

Montijn, J.S., Meijer, G.T., Lansink, C.S., Pennartz, C.M.A., 2016. Population-Level Neural Codes Are Robust to Single-Neuron Variability from a Multidimensional Coding Perspective. Cell Rep. 16, 2486–2498. https://doi.org/10.1016/j.celrep.2016.07.065

Montijn, J.S., Seignette, K., Howlett, M.H., Cazemier, J.L., Kamermans, M., Levelt, C.N., Heimel, J.A., 2021. A parameter-free statistical test for neuronal responsiveness. eLife 10, e71969. https://doi.org/10.7554/eLife.71969

Morrell, F., 1972. Visual System’s View of Acoustic Space. Nature 238, 44–46. https://doi.org/10.1038/238044a0

Musall, S., Kaufman, M.T., Juavinett, A.L., Gluf, S., Churchland, A.K., 2019. Single-trial neural dynamics are dominated by richly varied movements. Nat. Neurosci. 22, 1677–1686. https://doi.org/10.1038/s41593-019-0502-4

Niell, C.M., Stryker, M.P., 2010. Modulation of Visual Responses by Behavioral State in Mouse Visual Cortex. Neuron 65, 472– 479. https://doi.org/10.1016/j.neuron.2010.01.033

Niell, C.M., Stryker, M.P., 2008. Highly Selective Receptive Fields in Mouse Visual Cortex. J. Neurosci. Off. J. Soc. Neurosci. 28, 7520–7536. https://doi.org/10.1523/JNEUROSCI.0623-08.2008

Nikbakht, N., Tafreshiha, A., Zoccolan, D., Diamond, M.E., 2018. Supralinear and Supramodal Integration of Visual and Tactile Signals in Rats: Psychophysics and Neuronal Mechanisms. Neuron 97, 626–639. https://doi.org/10.1016/j.neuron.2018.01.003

Oh, S.W., Harris, J.A., Ng, L., Winslow, B., Cain, N., Mihalas, S., Wang, Q., Lau, C., Kuan, L., Henry, A.M., Mortrud, M.T., Ouellette, B., Nguyen, T.N., Sorensen, S.A., Slaughterbeck, C.R., Wakeman, W., Li, Y., Feng, D., Ho, A., Nicholas, E., Hirokawa, K.E., Bohn, P., Joines, K.M., Peng, H., Hawrylycz, M.J., Phillips, J.W., Hohmann, J.G., Wohnoutka, P., Gerfen, C.R., Koch, C., Bernard, A., Dang, C., Jones, A.R., Zeng, H., 2014. A mesoscale connectome of the mouse brain. Nature 508, 207–214. https://doi.org/10.1038/nature13186

Oude Lohuis, M.N., Pie, J.L., Marchesi, P., Montijn, J.S., de Kock, C.P.J., Pennartz, C.M.A., Olcese, U., 2022. Multisensory task demands temporally extend the causal requirement for visual cortex in perception. Nat. Commun. 13, 2864. https://doi.org/10.1038/s41467-022-30600-4

Paperna, T., Malach, R., 1991. Patterns of sensory intermodality relationships in the cerebral cortex of the rat. J. Comp. Neurol. 308, 432–456.

Pedregosa, F., Varoquaux, G., Gramfort, A., Michel, V., Thirion, B., Grisel, O., Blondel, M., Louppe, G., Prettenhofer, P., Weiss, R., Weiss, R.J., VanderPlas, J., Passos, A., Cournapeau, D., Brucher, M., Perrot, M., Duchesnay, E., 2011. Scikit-learn: Machine Learning in Python. J Mach Learn Res 12, 2825–2830.

Pennartz, C.M., 2015. The brain’s representational power: on consciousness and the integration of modalities. MIT Press.

Pennartz, C.M.A., 2009. Identification and integration of sensory modalities: Neural basis and relation to consciousness. Conscious. Cogn. 18, 718–739. https://doi.org/10.1016/j.concog.2009.03.003

Pennartz, C.M.A., Dora, S., Muckli, L., Lorteije, J.A.M., 2019. Towards a Unified View on Pathways and Functions of Neural Recurrent Processing. Trends Neurosci. 42, 589–603. https://doi.org/10.1016/j.tins.2019.07.005

Petro, L.S., Paton, A.T., Muckli, L., 2017. Contextual modulation of primary visual cortex by auditory signals. Philos. Trans. R. Soc. B Biol. Sci. 372, 20160104. https://doi.org/10.1098/rstb.2016.0104

Repp, B.H., Penel, A., 2002. Auditory dominance in temporal processing: New evidence from synchronization with simultaneous visual and auditory sequences. J. Exp. Psychol. Hum. Percept. Perform. 28, 1085–1099. https://doi.org/10.1037/0096-1523.28.5.1085

Ringach, D.L., Shapley, R.M., Hawken, M.J., 2002. Orientation Selectivity in Macaque V1: Diversity and Laminar Dependence. J. Neurosci. 22, 5639–5651. https://doi.org/10.1523/JNEUROSCI.22-13-05639.2002

Rockland, K.S., Ojima, H., 2003. Multisensory convergence in calcarine visual areas in macaque monkey. Int. J. Psychophysiol., Current findings in multisensory research 50, 19–26. https://doi.org/10.1016/S0167-8760(03)00121-1

Rossant, C., Kadir, S.N., Goodman, D.F.M., Schulman, J., Hunter, M.L.D., Saleem, A.B., Grosmark, A., Belluscio, M., Denfield, G.H., Ecker, A.S., Tolias, A.S., Solomon, S., Buzsáki, G., Carandini, M., Harris, K.D., 2016. Spike sorting for large, dense electrode arrays. Nat. Neurosci. 19, 634– 641. https://doi.org/10.1038/nn.4268

Runyan, C.A., Piasini, E., Panzeri, S., Harvey, C.D., 2017. Distinct timescales of population coding across cortex. Nature 548, 92–96. https://doi.org/10.1038/nature23020

Sakata, S., Harris, K.D., 2009. Laminar Structure of Spontaneous and Sensory-Evoked Population Activity in Auditory Cortex. Neuron 64, 404–418. https://doi.org/10.1016/j.neuron.2009.09.020

Schmitzer-Torbert, N., Jackson, J., Henze, D., Harris, K., Redish, A.D., 2005. Quantitative measures of cluster quality for use in extracellular recordings. Neuroscience 131, 1–11. https://doi.org/10.1016/j.neuroscience.2004.09.066

Schnabel, U.H., Bossens, C., Lorteije, J.A.M., Self, M.W., Op de Beeck, H., Roelfsema, P.R., 2018. Figure-ground perception in the awake mouse and neuronal activity elicited by figure-ground stimuli in primary visual cortex. Sci. Rep. 8. https://doi.org/10.1038/s41598-018-36087-8

Schneider, D.M., Nelson, A., Mooney, R., 2014. A synaptic and circuit basis for corollary discharge in the auditory cortex. Nature 513, 189–194. https://doi.org/10.1038/nature13724

Senzai, Y., Fernandez-Ruiz, A., Buzsáki, G., 2019. Layer-Specific Physiological Features and Interlaminar Interactions in the Primary Visual Cortex of the Mouse. Neuron. https://doi.org/10.1016/j.neuron.2018.12.009

Shams, L., Kamitani, Y., Shimojo, S., 2000. Illusions: What you see is what you hear. Nature 408, 788–788. https://doi.org/10.1038/35048669

Shepard, R.N., 1964. Circularity in Judgments of Relative Pitch. J. Acoust. Soc. Am. 36, 2346–2353. https://doi.org/10.1121/1.1919362

Sheppard, J.P., Raposo, D., Churchland, A.K., 2013. Dynamic weighting of multisensory stimuli shapes decision-making in rats and humans. J. Vis. 13, 4. https://doi.org/10.1167/13.6.4

Song, Y.-H., Kim, J.-H., Jeong, H.-W., Choi, I., Jeong, D., Kim, K., Lee, S.-H., 2017. A Neural Circuit for Auditory Dominance over Visual Perception. Neuron 93, 940–954. https://doi.org/10.1016/j.neuron.2017.01.006

Spinelli, D.N., Starr, A., Barrett, T.W., 1968. Auditory specificity in unit recordings from cat’s visual cortex. Exp. Neurol. 22, 75–84. https://doi.org/10.1016/0014-4886(68)90020-4

Spongr, V.P., Flood, D.G., Frisina, R.D., Salvi, R.J., 1997. Quantitative measures of hair cell loss in CBA and C57BL/6 mice throughout their life spans. J. Acoust. Soc. Am. 101, 3546–3553. https://doi.org/10.1121/1.418315

Stein, B.E., Stanford, T.R., 2008. Multisensory integration: current issues from the perspective of the single neuron. Nat. Rev. Neurosci. 9, 255–266. https://doi.org/10.1038/nrn2331

Steinmetz, N.A., Zatka-Haas, P., Carandini, M., Harris, K.D., 2019. Distributed coding of choice, action and engagement across the mouse brain. Nature 576, 266–273. https://doi.org/10.1038/s41586-019-1787-x

Stringer, C., Pachitariu, M., Steinmetz, N., Reddy, C.B., Carandini, M., Harris, K.D., 2019. Spontaneous behaviors drive multidimensional, brainwide activity. Science 364, 255– 255. https://doi.org/10.1126/science.aav7893

Vaknin, G., DiScenna, P.G., Teyler, T.J., 1988. A method for calculating current source density (CSD) analysis without resorting to recording sites outside the sampling volume. J. Neurosci. Methods 24, 131–135. https://doi.org/10.1016/0165-0270(88)90056-8

Van der Werf, Y.D., Witter, M.P., Groenewegen, H.J., 2002. The intralaminar and midline nuclei of the thalamus. Anatomical and functional evidence for participation in processes of arousal and awareness. Brain Res. Rev. 39, 107–140. https://doi.org/10.1016/S0165-0173(02)00181-9

Vinck, M., Batista-Brito, R., Knoblich, U., Cardin, J.A., 2015. Arousal and Locomotion Make Distinct Contributions to Cortical Activity Patterns and Visual Encoding. Neuron 86, 740–754. https://doi.org/10.1016/j.neuron.2015.03.028

Vinck, M., Bos, J.J., Van Mourik-Donga, L.A., Oplaat, K.T., Klein, G.A., Jackson, J.C., Gentet, L.J., Pennartz, C.M.A., 2016. Cell-Type and State-Dependent Synchronization among Rodent Somatosensory, Visual, Perirhinal Cortex, and Hippocampus CA1. Front. Syst. Neurosci. 9. https://doi.org/10.3389/fnsys.2015.00187

Williams, A.M., Angeloni, C.F., Geffen, M.N., 2021. Sound improves neuronal encoding of visual stimuli in mouse primary visual cortex. BioRxiv 2021.08.03.454738. https://doi.org/10.1101/2021.08.03.454738

Zagha, E., Erlich, J.C., Lee, S., Lur, G., O’Connor, D.H., Steinmetz, N.A., Stringer, C., Yang, H., 2022. The importance of accounting for movement when relating neuronal activity to sensory and cognitive processes. J. Neurosci. https://doi.org/10.1523/JNEUROSCI.1919-21.2021

